# The amplification of genetic factors for early vocabulary during children’s language and literacy development

**DOI:** 10.1101/863118

**Authors:** Ellen Verhoef, Chin Yang Shapland, Simon E. Fisher, Philip S. Dale, Beate St Pourcain

## Abstract

The heritability of language and literacy skills increases during development. The underlying mechanisms are little understood, and may involve (i) the amplification of early genetic influences and/or (ii) the emergence of novel genetic factors (innovation). Here, we use multivariate structural equation models to quantify these processes, as captured by genome-wide genetic markers. Studying expressive and receptive vocabulary at 38 months and subsequent language, literacy and cognitive skills (7-13 years) in unrelated children (ALSPAC: N≤6,092), we found little support for genetic innovation during mid-childhood and adolescence. Instead, genetic factors for early vocabulary, especially those unique to receptive skills, were amplified. Explaining as little as 3.9%(SE=1.8%) variation in early language, the same genetic influences accounted for 25.7%(SE=6.4%) to 45.1%(SE=7.6%) variation in verbal intelligence and literacy skills, but also performance intelligence, capturing the majority of SNP-heritability (≤99%). This suggests that complex verbal and non-verbal cognitive skills originate developmentaly in early receptive language.

## Introduction

Individual differences in vocabulary during the preschool period are predictive of many later language- and literacy-related skills^1–4^, an important component of academic achievement^5^. For example, a latent factor consisting of infant expressive and receptive vocabulary size at 16-24 months was found to predict vocabulary size, as well as performance on tests of phonological awareness, reading accuracy and reading comprehension in children five years later^3^. Similarly, infants with a larger expressive vocabulary at 24 months subsequently showed a larger vocabulary as well as better decoding, word recognition, and passage comprehension skills when assessed up to primary school^4^.

Associations between infant vocabulary and language and literacy skills during later life may arise due to shared underlying aetiologies. According to the “simple view of reading” theory, reading comprehension is the product of both printed word recognition (decoding) and oral language comprehension^6^. Early vocabulary is a central component of both these abilities^7^. Decoding is substantially based on phonological awareness (i.e. the awareness of sound structures of speech), which develops in the prescohol period and has been shown to be related to vocabulary size; listening comprehension (i.e. the understanding of spoken language), particularly bottom-up processing, necessarily begins with vocabulary comprehension. Spelling performance is also closely related to phonological awareness and other phonological abilities^8^. However, the biological processes that underlie these complex developmental interrelationships are only partially understood.

Variation in expressive and receptive vocabulary size, assessed during the first four years of life, is modestly heritable, while genetic influences on language and literacy skills assessed from mid-childhood to early adolescence are moderate to strong^9–12^. Specifically, longitudinal twin studies reported heritabilities (twin-h^2^) of 22%-28% for a combined language measure including expressive vocabulary at 2, 3 and 4 years of age^10^. Similar estimates were obtained for expressive vocabulary at 15-18 and 24-30 months of age in independent population-based samples, using genome-wide single-nucleotide polymorphism (SNP) information (SNP-h^2^=13%-14%)^11^. In contrast, the heritability for language and literacy skills assessed from mid-childhood onwards is larger, with twin-h^2^ estimates of 47%-72%^9,10^ and SNP-h^2^ estimates of 32%-54%^12^. However, developmental stages nonetheless genetically overlap, as shown by moderate genetic correlations reported in twin research^9,10^.

The increase in heritability from early childhood to adolescence has been reported for many cognitive skills^13,14^, suggesting overarching aetiological mechanisms that may involve processes of genetic innovation and amplification^15^. Innovation refers to novel genetic factors emerging during development (i.e. previously unrelated genetic variation becoming associated with a trait). In contrast, amplification refers to genetically stable influences that are active throughout development, explaining increasingly more variation with progressing age^13^. A meta-analysis of twin studies on cognitive abilities suggested that novel genetic influences predominate during the transition from early to middle childhood, but wane quickly, with enhanced genetic stability and amplification processes dominating from 8 years of age onwards^13^. This developmental paradigm is consistent with twin study findings examining genetic links between early language (including expressive vocabulary and syntax skills between 2-4 years of age) and mid-childhood/adolescent language^10^ and reading^9^, based on latent factor models. Thus, it is possible that innovation rather than amplification processes will account for the observed increase in heritability during language and literacy development, not only in twins^9,10^, but in all typically developing children. Furthermore, these processes may represent a developmental paradigm that has relevance not only for language and literacy skills, but cognitive functioning in general, possibly involving “generalist genes” that impact on many related traits^16^.

However, beyond latent factor twin analyses^9,10^, the specific processes genetically linking early vocabulary skills with language, literacy and cognition later in life are little characterised. In particular, genetic relationships with early receptive vocabulary are unknown, and the spectrum of interrelated skills, shaping language, literacy and cognition, affected by amplification processes is only partially understood. Here, we use SNP information from directly genotyped common genetic markers and structural equation models to quantify these genetic mechanisms within a sample of unrelated children from the Avon Longitudinal Study of Parents And Children (ALSPAC, N≤6,092). Specifically, we study expressive and receptive vocabulary at 38 months and a wide range of later language- and literacy-related skills (7-13 years, including reading, spelling, phonemic awareness, listening comprehension, non-word repetition) as well as verbal and non-verbal intelligence scores, seeking evidence for innovation and/or amplification processes.

## Methods

### Participants

All participants were drawn from ALSPAC, a UK population-based longitudinal pregnancy-ascertained birth cohort (estimated birth date: 1991-1992, Supplementary Methods)^17,18^. The ALSPAC Ethics and Law Committee and the Local Research Ethics Committees provided ethical approval for the study. Consent for biological samples has been collected in accordance with the Human Tissue Act (2004). Informed consent for the use of data collected via questionnaires and clinics was obtained from participants following the recommendations of the ALSPAC Ethics and Law Committee at the time.

ALSPAC participants were genotyped using the Illumina HumanHap550 quad chip genotyping platforms. Standard genomic quality control was performed at both the SNP and individual level using PLINK (v1.07)^19^ (Supplementary Methods). After quality control, 8,226 children and 465,740 SNPs remained.

### Measures

#### Early-life vocabulary

Expressive and receptive vocabulary were assessed at 38 months using parent reports and age-specific defined word lists adapted from the MacArthur Communicative Development Inventory Words & Sentences (CDI)^20^. Parents were asked whether their child was able to say, understand or both say and understand a word from a list of 123 words. Expressive vocabulary was defined as the number of words a child was able to say or say and understand, whereas receptive vocabulary was defined as the number of words a child could understand or say and understand. In total, 6,092 children had both phenotypic and genome-wide genetic data available (Table 1).

**Table 1:**
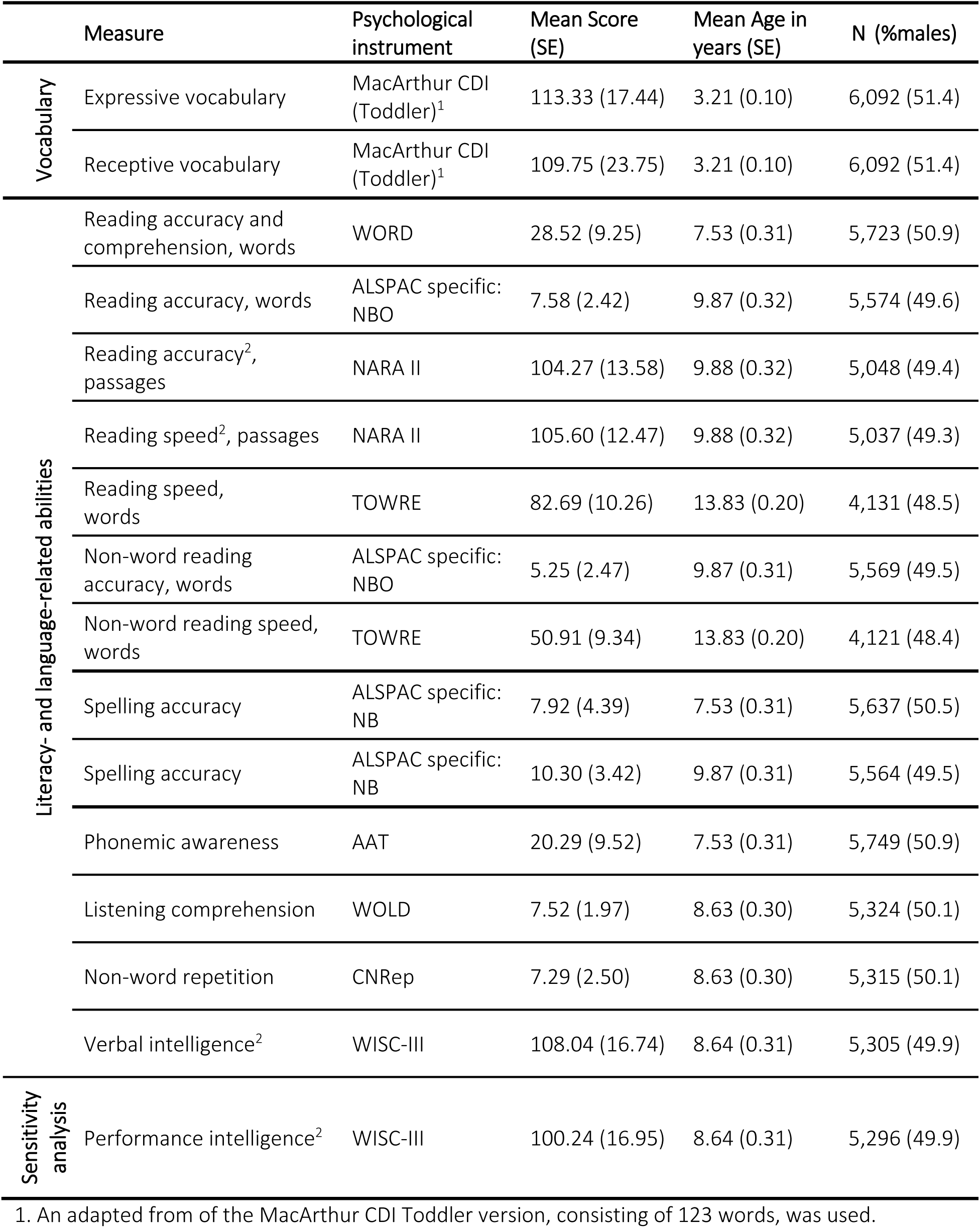
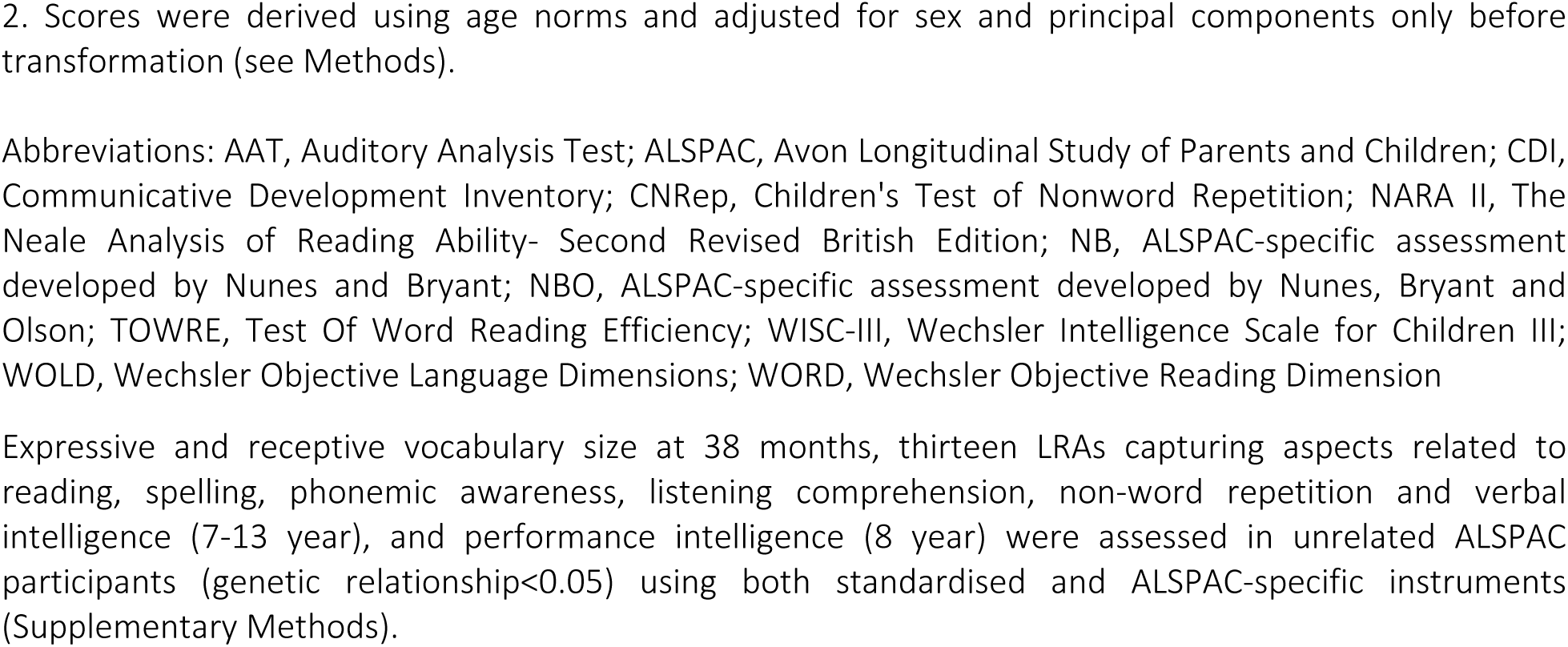
Early vocabulary and mid-childhood to adolescence literacy, language and cognition abilities in the Avon Longitudinal Study of Parents and Children

|  | Measure | Psychological instrument | Mean Score (SE) | Mean Age in years (SE) | N (%males) |
| --- | --- | --- | --- | --- | --- |
| Vocabulary | Expressive vocabulary | MacArthur CDI (Toddler)<sup>1</sup> | 113.33 (17.44) | 3.21 (0.10) | 6,092 (51.4) |
|  | Receptive vocabulary | MacArthur CDI (Toddler)<sup>1</sup> | 109.75 (23.75) | 3.21 (0.10) | 6,092 (51.4) |
| Literacy- and language-related abilities | Reading accuracy and comprehension, words | WORD | 28.52 (9.25) | 7.53 (0.31) | 5,723 (50.9) |
|  | Reading accuracy, words | ALSPAC specific: NBO | 7.58 (2.42) | 9.87 (0.32) | 5,574 (49.6) |
|  | Reading accuracy<sup>2</sup>, passages | NARA II | 104.27 (13.58) | 9.88 (0.32) | 5,048 (49.4) |
|  | Reading speed<sup>2</sup>, passages | NARA II | 105.60 (12.47) | 9.88 (0.32) | 5,037 (49.3) |
|  | Reading speed, words | TOWRE | 82.69 (10.26) | 13.83 (0.20) | 4,131 (48.5) |
|  | Non-word reading accuracy, words | ALSPAC specific: NBO | 5.25 (2.47) | 9.87 (0.31) | 5,569 (49.5) |
|  | Non-word reading speed, words | TOWRE | 50.91 (9.34) | 13.83 (0.20) | 4,121 (48.4) |
|  | Spelling accuracy | ALSPAC specific: NB | 7.92 (4.39) | 7.53 (0.31) | 5,637 (50.5) |
|  | Spelling accuracy | ALSPAC specific: NB | 10.30 (3.42) | 9.87 (0.31) | 5,564 (49.5) |
|  | Phonemic awareness | AAT | 20.29 (9.52) | 7.53 (0.31) | 5,749 (50.9) |
|  | Listening comprehension | WOLD | 7.52 (1.97) | 8.63 (0.30) | 5,324 (50.1) |
|  | Non-word repetition | CNRep | 7.29 (2.50) | 8.63 (0.30) | 5,315 (50.1) |
|  | Verbal intelligence<sup>2</sup> | WISC-III | 108.04 (16.74) | 8.64 (0.31) | 5,305 (49.9) |
| Sensitivity analysis | Performance intelligence<sup>2</sup> | WISC-III | 100.24 (16.95) | 8.64 (0.31) | 5,296 (49.9) |

1. An adapted from of the MacArthur CDI Toddler version, consisting of 123 words, was used.
2. Scores were derived using age norms and adjusted for sex and principal components only before transformation (see Methods).
Abbreviations: AAT, Auditory Analysis Test; ALSPAC, Avon Longitudinal Study of Parents and Children; CDI, Communicative Development Inventory; CNRep, Children's Test of Nonword Repetition; NARA II, The Neale Analysis of Reading Ability- Second Revised British Edition; NB, ALSPAC-specific assessment developed by Nunes and Bryant; NBO, ALSPAC-specific assessment developed by Nunes, Bryant and Olson; TOWRE, Test Of Word Reading Efficiency; WISC-III, Wechsler Intelligence Scale for Children III; WOLD, Wechsler Objective Language Dimensions; WORD, Wechsler Objective Reading Dimension
Expressive and receptive vocabulary size at 38 months, thirteen LRAs capturing aspects related to reading, spelling, phonemic awareness, listening comprehension, non-word repetition and verbal intelligence (7-13 year), and performance intelligence (8 year) were assessed in unrelated ALSPAC participants (genetic relationship<0.05) using both standardised and ALSPAC-specific instruments (Supplementary Methods).

#### Mid-childhood/adolescent language- and literacy-related abilities

Thirteen measures capturing reading, spelling, phonemic awareness, listening comprehension, non-word repetition and verbal intelligence were assessed from mid-childhood onwards (7-13 year, N≤5,749) using both standardised and ALSPAC-specific instruments (Table 1, Supplementary Methods). Combined word reading accuracy and comprehension (age 7 years) was measured using the basic reading subtest of the Wechsler Objective Reading Dimensions (WORD)^21^ assessment in addition to word and non-word reading accuracy scores (age 9 years) using an ALSPAC-specific measure^22^. Passage reading accuracy and speed (age 9 years) was captured with the revised Neale Analysis of Reading Ability (NARA II)^23^ and word and non-word reading speed (age 13 years) with the Test of Word Reading Efficiency (TOWRE)^24^. Spelling accuracy (age 7 and 9 years) was assessed with an ALSPAC-specific measure (Supplementary Methods). Phonemic awareness (age 7 years) was measured with the Auditory Analysis Test (AAT)^25^ and listening comprehension, non-word repetition and verbal intelligence quotient (VIQ) scores (all age 8 years) were assessed with a subset of the Wechsler Objective Language Dimensions (WOLD)^26^ test, an adaptation of the Children’s Test of Nonword Repetition (CNRep)^27^ and the Wechsler Intelligence Scale for Children (WISC-III)^28^ respectively.

#### Mid-childhood performance intelligence

For sensitivity analyses, we studied performance intelligence quotient (PIQ) scores as assessed using the WISC-III^28^ (Table 1, Supplementary Methods).

#### Phenotype transformation

Vocabulary, LRA and PIQ scores were rank-transformed to achieve normality and to allow for comparisons of genetic effects across different psychological instruments. Vocabulary measures were residualised for sex, age, age^2^ and the two most significant ancestry-informative principal components, calculated using EIGENSOFT^29^ (v6.1.4). LRAs and PIQ measures were residualised for sex, age (unless measures were derived using age-specific norms) and the two most significant ancestry-informative principal components.

### Analysest

#### Phenotypic correlations

Phenotypic correlations (r_p_) were calculated for untransformed and rank-transformed scores using Spearman rank-correlation and Pearson correlation coefficients respectively. Patterns were highly similar for untransformed and transformed scores (Supplementary Figure 1-2).

#### Genome-wide Complex Trait Analysis

SNP-h^2^ was estimated using Restricted Maximum Likelihood (REML) analyses as implemented in Genome-wide Complex Trait Analysis (GCTA, v1.26.0) software^30^. This method examines unrelated individuals pair by pair and correlates their genetic similarity with their phenotypic similarity. Genetic similarity can be captured in a genetic-relationship matrix (GRM)^30^, a matrix with as many columns and rows as individuals, that was created using PLINK (v1.9)^19^. GRMs were constructed for individuals with a genetic relationship <0.05 (N_individuals_≤6,092) and based on directly genotyped SNPs only. (N_SNPs_=465,740). Genetic correlations (r_g_), reflecting the extent to which two measures are influenced by the same genetic factors, were estimated using bivariate REML^31^ within GCTA and the GRM as described above.

#### Multivariate genetic analyses

To quantify shared and unique genetic influences contributing to vocabulary at 38 months and mid-childhood/adolescent LRAs, we studied their genetic variance/co-variance with multivariate Genetic-relationship-matrix Structural Equation Modelling (GSEM) techniques^32^. The derived path models are analogous to twin research methodologies^33,34^. However, like GCTA, they use GRMs to estimate genetic variances and covariance structures between unrelated individuals^32^ (Supplementary Methods). Multivariate trait variances were modelled using a Cholesky decomposition^34^. This saturated model involves the decomposition of phenotypic variances into an ordered series of latent genetic and residual factors, as many as there are observed variables^34^ (Supplementary Methods). GSEM models were fitted using all available observations for children across development (R:gsem library, version 0.1.5). In addition to estimating path coefficients, we also utilised GSEM to estimate SNP-h^2^, genetic correlations, factorial co-heritability (the proportion of total genetic variance explained by a specific genetic factor) and bivariate heritability (the contribution of genetic factors to the observed phenotypic correlation between two measures)(Supplementary Methods).

Our data analysis strategy followed a two-step procedure: First, using GSEM, we fitted 13 trivariate structural equation models (SEMs), each consisting of expressive and receptive vocabulary at 38 months and one of the 13 LRAs (in this order, termed “forward” GSEM, Supplementary Figure 4a). Second, we carried out a meta-analysis of absolute GSEM path coefficients for these 13 models across pre-defined domains including (1) reading-related measures, (2) spelling-related measures, and (3) all LRA outcomes (Supplementary Table 2). Estimates were combined using random-effects meta-regression intercepts, accounting for interrelatedness between LRAs (R:metafor library, Rv3.2.0)^35^. For this, a variance/covariance matrix across measures was approximated by including the observed phenotypic correlation matrix, weighted by the standard errors of the path coefficients as estimated by GSEM, analogous to models accounting for correlated phylogenetic histories^36^. As part of sensitivity analyses, the order of the two vocabulary measures at 38 months was reversed within the 13 SEMs (termed “reverse” GSEM, Supplementary Figure 6a). To compare LRA genetic covariance patterns with non-verbal cognitive abilities, we also studied expressive and receptive vocabulary at 38 months together with PIQ at 8 years.

#### Experiment-wide significance threshold

The effective number of phenotypes was calculated based on phenotypic correlations using matrix Spectral Decomposition (matSpD)^37^, resulting in nine independent measures. This corresponds to an experiment-wide significance threshold of 0.005 (0.05/9).

## Results

### Phenotypic and genetic descriptives

Expressive and receptive vocabulary assessed at 38 months are modestly heritable as tagged by common genotyping information, with GCTA-SNP-h^2^ estimates of 18% (SNP-h^2^: 0.18(SE=0.06)) and 12% (SNP-h^2^: 0.12(SE=0.06)) respectively (Figure 1, Supplementary Table 1). GCTA-SNP-h^2^ estimates for LRAs assessed during mid-childhood and adolescence, including reading abilities (comprehension, accuracy and speed), spelling abilities (accuracy), phonemic awareness, listening comprehension, non-word repetition and VIQ were moderate, reaching up to 54% (SNP-h^2^:0.54(SE=0.07)) (Figure 1, Supplementary Table 1), as previously reported^12^.

**Figure 1:**
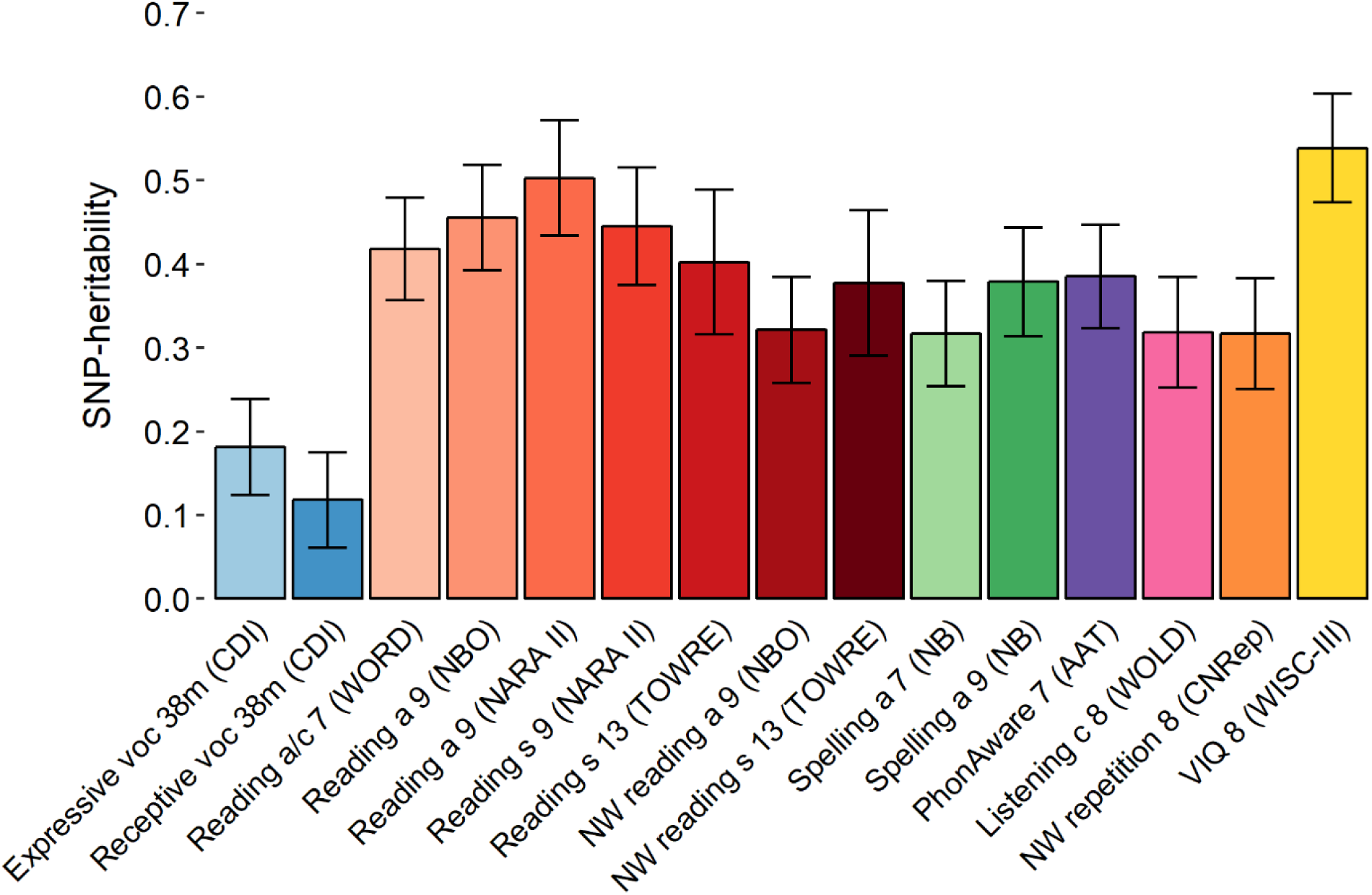
SNP-heritability estimates for early vocabulary and mid-childhood to adolescence literacy- and language-related abilities. Abbreviations: a, accuracy; AAT, Auditory Analysis Test; c, comprehension; CDI, Communicative Development Inventory; CNRep, Children’s Test of Nonword Repetition; m, months; NARA II, The Neale Analysis of Reading Ability-Second Revised British Edition; NB, ALSPAC-specific assessment developed by Nunes and Bryant; NBO, ALSPAC-specific assessment developed by Nunes, Bryant and Olson; NW, nonword; PhonAware, phonemic awareness; s, speed; TOWRE, Test Of Word Reading Efficiency; VIQ, verbal intelligence quotient; voc, vocabulary; WISC-III, Wechsler Intelligence Scale for Children III; WOLD, Wechsler Objective Language Dimensions; WORD, Wechsler Objective Reading Dimension SNP-heritability was estimated for each measure based on rank-transformed scores using Restricted Maximum Likelihood (REML) analyses as implemented in genome-wide complex trait analysis (GCTA) software. Analyses were based on independent individuals (genetic relationship of <0.05) and directly genotyped SNPs. Bars represent standard errors.

Consistent with phenotypic correlations between expressive and receptive vocabulary at 38 months (r_p_=0.63, Supplementary Figure 2), bivariate genetic correlations were strong (r_g_=0.86(SE=0.15), *P*=0.004, Supplementary Figure 3) and shared genetic influences accounted for ∼20% of the observed phenotypic overlap (bivariate heritability: 0.19(SE=0.07)). Both expressive and receptive vocabulary at 38 months were phenotypically also correlated with language and literacy skills later in life (Supplementary Figure 2). Phenotypic correlations of LRAs with receptive vocabulary ranged between 0.14 and 0.26, and with expressive vocabulary between 0.12 and 0.18 (Supplementary Figure 2). At the genetic level, receptive vocabulary was moderately to strongly linked with the entire spectrum of LRAs, with genetic correlations ranging from 0.58 (SE=0.21, *P*=0.001) to 0.95 (SE=0.23, *P*=1×10^−8^)(Supplementary Figure 3). Expressive vocabulary was genetically correlated only with VIQ scores at 8 years (r_g_=0.38(SE=0.14), *P*=0.003, Supplementary Figure 3).

### Structural equation modelling

Next, we modelled multivariate genetic variances between expressive and receptive vocabulary at 38 months and, in turn, each of the 13 mid-childhood/adolescent LRAs using GSEM. Within each forward GSEM model, the estimated path coefficients link to shared and unique genetic variance components through structural equations (Supplementary Methods). SNP-h^2^ estimates were consistent between GCTA and GSEM for all SEMs studied (Supplementary Table 1).

Squared path coefficients for the first genetic factor (A1) fully explain genetic variance in expressive vocabulary at 38 months (a_11_) and genetic variance that is shared with receptive vocabulary (a_21_) and the selected LRA (a_31_, Supplementary Figure 4a). The first two path coefficients (a_11,_ a_21_) were nearly identical across all 13 models (Supplementary Figure 4), and are reported here for the model including expressive and receptive vocabulary at 38 months and VIQ scores at 8 years (Figure 2a). According to this model, the first genetic factor explained 17.7%(SE=5.7%) of the phenotypic variance in expressive vocabulary (path-coefficient a_11_=-0.42(SE=0.06), *P*=4×10^−10^, Figure 2a-b), corresponding to the SNP-h^2^ of the trait. It also accounted for 9.0%(SE=4.5%) of the phenotypic variance in receptive vocabulary (path-coefficient a_21_=-0.30(SE=0.08), *P*=7×10^−5^, Figure 2a-b), capturing approximately two thirds of its total genetic variance (factorial co-heritability: 0.67(SE=0.17)). In addition, this first genetic factor explained 8.6%(SE=6.0%) of the phenotypic variance in VIQ scores at 8 years (path-coefficient a_31_=-0.29(SE=0.10), *P*=0.004, Figure 2a-b), but not other LRAs (Supplementary Figure 4). This corresponds to ∼16% of the total genetic variance in VIQ at 8 years (factorial co-heritability: 0.16(SE=0.11), Supplementary Table 3).

**Figure 2:**
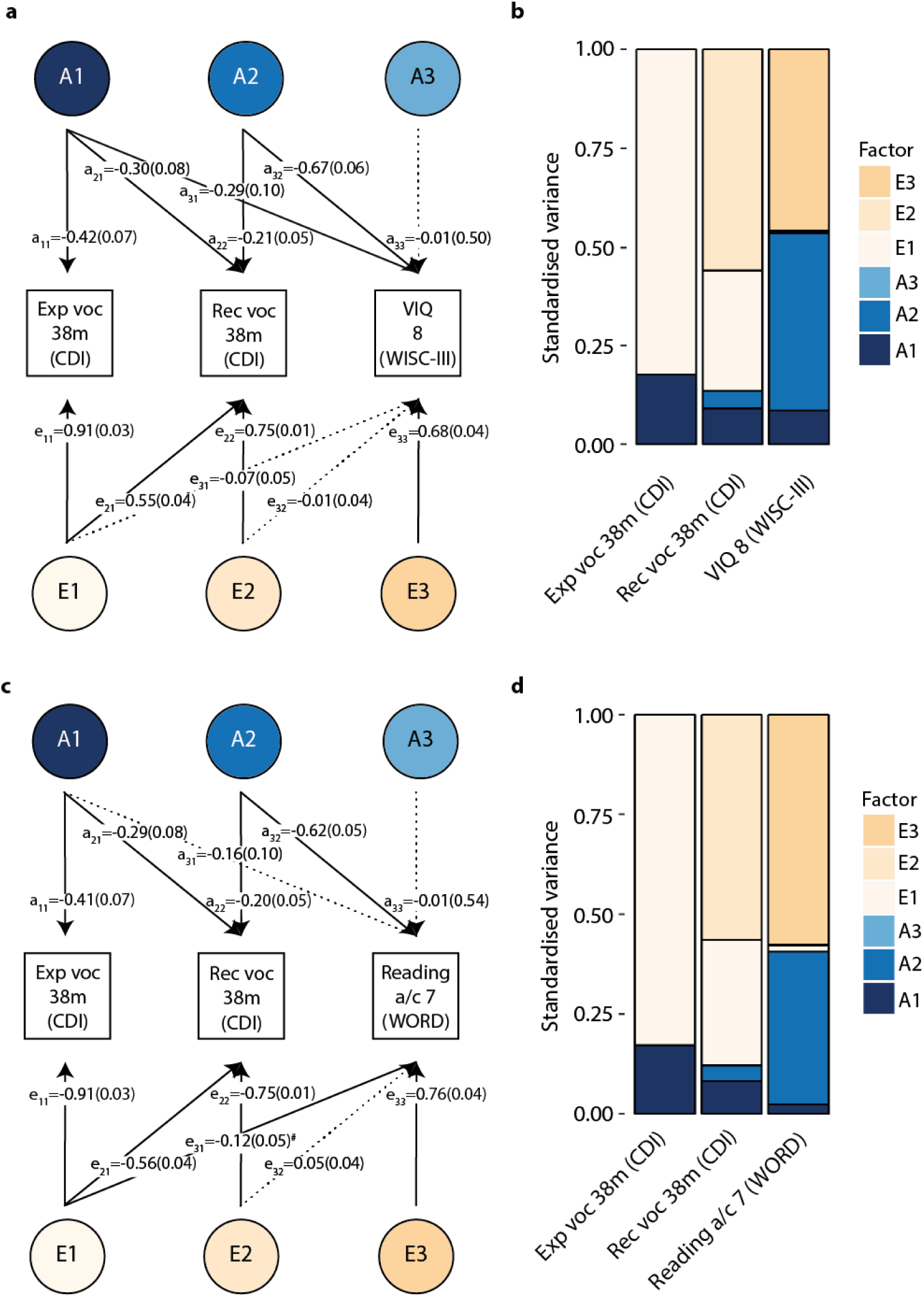
Path models and variance plots for early vocabulary and verbal intelligence or reading accuracy/comprehension in mid-childhood. Abbreviations: a, accuracy; c, comprehension; CDI, Communicative Development Inventory; Exp, expressive; m, months; Rec, receptive, Voc, vocabulary; VIQ, verbal intelligence quotient; WISC-III, Wechsler Intelligence Scale for Children III; WORD, Wechsler Objective Reading Dimension # Path coefficient passing nominal significance (*P*≤0.05), but not the experiment-wide significance threshold (*P*≤0.005). Cholesky decompositions were fitted using GSEM, according to forward GSEMs and based on all available observations for children across development (N≤6,092). **(a,c)** Path models of standardised path coefficients and corresponding standard errors for Cholesky decompositions of vocabulary at 38 months and **(a)** verbal intelligence assessed at 8 years or **(c)** reading accuracy/comprehension at 7 years. Solid lines indicate path coefficients passing a *P*-value threshold of *P*≤0.05, dashed lines indicate non-significant path coefficients (*P*>0.05). **(b,d)** Standardised variance explained by genetic and residual factors modelled in a,c using Cholesky decompositions of vocabulary at 38 months and **(b)** verbal intelligence assessed at 8 years or **(d)** reading accuracy/comprehension assessed at 7 years.

Squared path coefficients for the second genetic factor (A2) capture unique genetic variance in receptive vocabulary at 38 months, independent of expressive vocabulary (a_22_, Supplementary Figure 4a), and the extent to which this genetic variance is shared with the selected LRA (a_32_, Supplementary Figure 4a). Nearly identically across the 13 GSEM models, the second genetic factor described a further 4.5% (SE=2.0%) of the phenotypic variance of receptive vocabulary at 38 months (path coefficient a_22_=-0.21(SE=0.05), *P*=4×10^−6^, here shown for the model including VIQ, Figure 2a-b). Thus, about a third of the genetic variance in receptive vocabulary is unique and not shared with expressive vocabulary at the same age (factorial co-heritability=0.33(SE=0.17)). Importantly, this small proportion of genetic variance was amplified and accounted for the majority of genetic influences contributing to subsequent VIQ, reading and spelling abilities (path-coefficient a_32_, Figure 2, Supplementary Figure 4-5, Supplementary Table 3). For example, this genetic factor accounted for 45.1%(SE=7.6%) of the phenotypic variance in VIQ scores at 8 years (path-coefficient a_32_=-0.67(SE=0.06), *P*<1×10^−10^, Figure 2a-b), corresponding to >80% of the SNP-h^2^ (factorial co-heritability: 0.84(SE=0.11), Supplementary Table 3). Similarly, for literacy-related traits, the second genetic factor explained 38.2%(SE=6.0%) of the phenotypic variance in reading accuracy/comprehension at 7 years of age (path-coefficient a_32_=-0.62(SE=0.05), *P*<1×10^−10^, Figure 2c-d), entailing nearly the entire SNP-h^2^ of the measure (factorial co-heritability: 0.94(SE=0.08), Supplementary Table 3). Comparable patterns were observed for reading accuracy at 9 years (assessed with NARA II), reading speed at 9 years, reading and non-word reading speed at 13 years and spelling accuracy at 7 years, with ≥29.4% of phenotypic variation explained by genetic variance unique for receptive vocabulary (Supplementary Figure 4-5).

Squared path coefficients for the third genetic factor (A3) account for unique genetic variance in the studied LRAs, independent of genetic factors contributing to both expressive and receptive vocabulary at 38 months (a_33_, Supplementary Figure 4a). Contrary to our initial hypothesis, we found little evidence for novel genetic LRA influences arising after early childhood (Figure 2, Supplementary Figure 4).

A meta-analysis of absolute Cholesky path coefficients across all 13 SEM models (Supplementary Table 2), correcting for phenotypic inter-correlations (Supplementary Figure 2), confirmed the amplification of genetic influences that are unique to receptive vocabulary at 38 months (meta-path-coefficient a_32_=0.62(0.06), *P*<1×10^−10^, Table 2). Nominal evidence was also found for the amplification of genetic influences that capture the entirety of expressive vocabulary at 38 months (meta-path-coefficient a_31_=0.20(SE=0.08), *P*=0.009, Table 2), although it did not pass the experiment-wide multiple testing threshold. Consistent with individual GSEM models, there was little meta-analytic evidence for novel genetic influences arising after early childhood (meta-path-coefficient a_33_=0.34(SE=0.29), *P*=0.24, Table 2). Literacy-specific meta-analyses of reading measures only and spelling measures only, suggested that developmental genetic amplification patterns involve primarily, but not exclusively, reading-related abilities (Table 2).

**Table 2:** Meta-analysis across pre-defined language- and literacy-related ability combinations.

| Path |  | All LRAs (N=13) |  |  | Reading (N=7) |  |  | Spelling (N=2) |  |  |
| --- | --- | --- | --- | --- | --- | --- | --- | --- | --- | --- |
|  |  | Coefficient (SE) | *P* | *P*<sub>het</sub> | Coefficient (SE) | *P* | *P*<sub>het</sub> | Coefficient (SE) | *P* | *P*<sub>het</sub> |
| Genetic influences | a<sub>11</sub> | 0.42(0.05) | <1x10<sup>-10</sup> | 1.00 | 0.42(0.06) | <1x10<sup>-10</sup> | 1.00 | 0.42(0.06) | <1x10<sup>-10</sup> | 0.97 |
|  | a<sub>21</sub> | 0.29(0.06) | 1x10<sup>-6</sup> | 1.00 | 0.29(0.07) | 2x10<sup>-5</sup> | 1.00 | 0.29(0.08) | 2x10<sup>-4</sup> | 0.98 |
|  | a<sub>31</sub> | 0.20(0.08) | 9x10<sup>-3</sup> | 0.70 | 0.19(0.09) | 0.03 | 0.70 | 0.19(0.10) | 0.04 | 0.54 |
|  | a<sub>22</sub> | 0.19(0.05) | 3x10<sup>-5</sup> | 1.00 | 0.19(0.05) | 3x10<sup>-5</sup> | 1.00 | 0.17(0.07) | 8x10<sup>-3</sup> | 0.99 |
|  | a<sub>32</sub> | 0.62(0.06) | <1x10<sup>-10</sup> | 0.97 | 0.62(0.06) | <1x10<sup>-10</sup> | 0.94 | 0.52(0.13) | 1x10<sup>-4</sup> | 0.50 |
|  | a<sub>33</sub> | 0.34(0.29) | 0.24 | 1.00 | 0.37(0.29) | 0.20 | 0.95 | 0.38(0.32) | 0.23 | 0.65 |
| Residual influences | e<sub>11</sub> | 0.91(0.02) | <1x10<sup>-10</sup> | 1.00 | 0.91(0.03) | <1x10<sup>-10</sup> | 1.00 | 0.91(0.03) | <1x10<sup>-10</sup> | 0.97 |
|  | e<sub>21</sub> | 0.56(0.03) | <1x10<sup>-10</sup> | 1.00 | 0.56(0.03) | <1x10<sup>-10</sup> | 1.00 | 0.56(0.04) | <1x10<sup>-10</sup> | 0.98 |
|  | e<sub>31</sub> | 0.08(0.04) | 0.02 | 0.62 | 0.09(0.04) | 0.03 | 0.25 | 0.07(0.04) | 0.11 | 0.98 |
|  | e<sub>22</sub> | 0.75(0.01) | <1x10<sup>-10</sup> | 1.00 | 0.75(0.01) | <1x10<sup>-10</sup> | 1.00 | 0.76(0.02) | <1x10<sup>-10</sup> | 0.99 |
|  | e<sub>32</sub> | 0.03(0.03) | 0.36 | 0.91 | 0.04(0.04) | 0.23 | 0.83 | 0.02(0.04) | 0.67 | 0.58 |
|  | e<sub>33</sub> | 0.78(0.03) | <1x10<sup>-10</sup> | 8x10<sup>-4</sup> | 0.76(0.04) | <1x10<sup>-10</sup> | 0.09 | 0.80(0.04) | <1x10<sup>-10</sup> | 0.25 |

Abbreviations: LRAs, literacy- and language-related abilities; *P*<sub>het</sub>, *P*-value for the test of heterogeneity
Absolute path coefficients for 13 structural equation models (forward GSEM) were meta-analysed across pre-defined domains accounting for interrelatedness between traits. Path coefficients were considered significant if they passed the experiment-wide *P*-value threshold of 0.005. Heterogeneity among effect estimates was assessed using Cochran's Q-test.

Variance decompositions using a Cholesky model are sensitive to the order that traits are incorporated into the model, although SNP-h^2^ estimations remain unchanged. We therefore created 13 additional GSEM models, reversing the order of expressive and receptive vocabulary at 38 months (reverse GSEM models, with path coefficients as detailed in Supplementary Figure 6a). Consistent with forward GSEM models, there was little evidence for novel LRA-related genetic factors emerging after early childhood (Supplementary Figure 6-7), as estimated with the third genetic factor (A3). However, for reverse GSEM, the first genetic factor (A1) captures the entire genetic variance of receptive vocabulary and to what extent it is shared with expressive vocabulary and later LRAs. This factor accounted for two thirds of the total genetic variance in expressive vocabulary (factorial co-heritability: 0.67(SE=0.17)), corresponding to 11.8%(SE=5.5%) of the phenotypic variance (shown for the GSEM model including VIQ). The second genetic factor (A2) captures genetic influences that are unique to expressive vocabulary (i.e. independent of receptive vocabulary) and explained an additional 5.9%(SE=3.0%) of its phenotypic and a third of its genetic variance (factorial co-heritability: 0.33(SE=0.17)). Both early genetic factors accounted for phenotypic variation in VIQ, reading and spelling abilities, but also phonemic awareness and/or non-word repetition (Supplementary Figure 6-7).

To identify the most predictive genetic variance components of early vocabulary using either forward or reverse GSEM models, we studied model-specific factorial co-heritabilities and bivariate heritabilities (which are identical for forward and reverse GSEM). The largest contribution to genetic variance in later LRAs was confirmed for genetic influences uniquely related to receptive vocabulary (A2, forward GSEM, Supplementary Figure 4a), explaining up to 95%(SE=20%) in LRA SNP-h^2^, especially for reading and VIQ scores (Supplementary Table 3). In comparison, the contribution of receptive vocabulary-related genetic influences that are shared with expressive vocabulary (A1, reverse GSEM, Supplementary Figure 6a) were lower, explaining only up to 73%(SE=20%) of LRA SNP-h^2^, although 95% confidence intervals overlap. For example, uniquely receptive vocabulary-related genetic influences explained 94%(SE=8%) of genetic variance for reading accuracy and comprehension at age 7, while receptive vocabulary-related genetic influences that are shared with expressive vocabulary explained only 57%(SE=23%) and did not pass the experiment-wide threshold (Supplementary Table 3). Consistently, genetic covariance between receptive vocabulary and later LRAs explained the majority of their phenotypic covariance, with bivariate heritability estimates of up to 1.00(SE=0.22), except for listening comprehension and non-word reading accuracy (Figure 3, Supplementary Table 4). In contrast, there was little evidence that genetic factors underlying expressive vocabulary, irrespective of its decomposition, substantially accounted for variation in LRAs (Figure 3, Supplementary Table 4), except for VIQ scores (0.69(SE=0.24)). Thus, the majority of genetic variation in later LRAs can be attributed to a small proportion of genetic variance in early language that uniquely captures receptive vocabulary and has been amplified during development.

**Figure 3:**
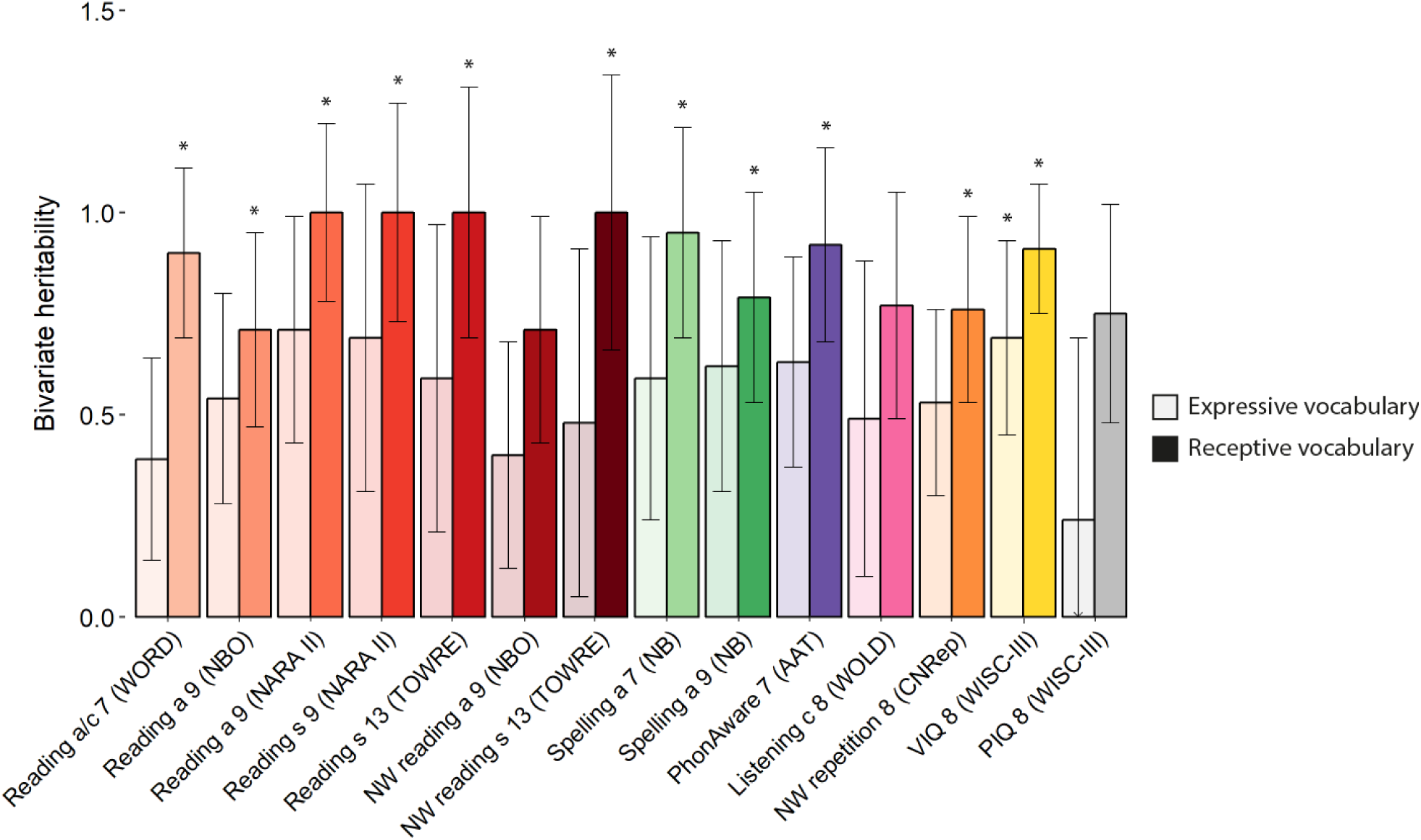
Bivariate heritability estimates. Abbreviations: a, accuracy; AAT, Auditory Analysis Test; c, comprehension; CNRep, Children’s Test of Nonword Repetition; NARA II, The Neale Analysis of Reading Ability-Second Revised British Edition; NB, ALSPAC-specific assessment developed by Nunes and Bryant; NBO, ALSPAC-specific assessment developed by Nunes, Bryant and Olson; NW, nonword; PhonAware, phonemic awareness; PIQ; performance intelligence quotient; s, speed; TOWRE, Test Of Word Reading Efficiency; VIQ, verbal intelligence quotient; WISC-III, Wechsler Intelligence Scale for Children III; WOLD, Wechsler Objective Language Dimensions; WORD, Wechsler Objective Reading Dimension * Bivariate heritability estimate passing the experiment-wide significance threshold (*P*≤0.005) Bivariate heritability estimates, reflecting the proportion of the phenotypic covariance that is accounted for by the genetic covariance. SEs were approximated by the SE of the genetic covariance divided by the phenotypic covariance (as the SE of the phenotypic covariance is small) and *P*-values are based on a Wald-test, assuming normality. Estimates are based on forward GSEM models, and reverse GSEM models provided nearly identical results (data not shown). Bivariate heritability estimates were truncated at one for reading a 9 (NARA II), reading s 9 (NARA II), reading s 13 (TOWRE) and NW reading s 13 (TOWRE). Bars represent standard errors.

Finally, we assessed whether the amplification of a genetic factor that is unique to receptive vocabulary extends to non-verbal cognitive tasks. For this, we studied GSEM models including expressive and receptive vocabulary at 38 months as well as PIQ at 8 years (Supplementary Figure 8). Findings were highly similar to results observed for VIQ and reading. More specifically, using forward GSEM, there was (i) a link between genetic influences unique to receptive vocabulary (A2) and PIQ, explaining 25.7%(SE=6.4%) of phenotypic variance in PIQ (path-coefficient a_32_=-0.51(SE=0.06), *P*<1×10^−10^, Supplementary Figure 8a-b; factorial co-heritability: 0.99(SE=0.04), Supplementary Table 3); and (ii) no support for genetic influences that are specific to PIQ and arise during mid-childhood (A3, Supplementary Figure 8a-b). Findings using reverse GSEM for PIQ were also similar to patterns observed for other LRAs (Supplementary Figure 6-7, 8c-d). However, evidence for the contribution of genetic factors to phenotypic covariance between receptive vocabulary and PIQ was less strong compared to literacy and verbal cognition skills and did not pass the experiment-wide threshold (bivariate heritability: 0.75(0.27), Figure 3, Supplementary Table 4), irrespective of forward or reverse GSEM.

## Discussion

This study provides evidence that the amplification of early vocabulary-related genetic factors plays a major role during later language and literacy development. Multivariate variance analyses using genome-wide data showed that genetic influences underlying receptive vocabulary at 38 months, and to a lesser extent expressive vocabulary at the same age, could fully account for genetic variation in many reading and spelling skills, but also verbal and non-verbal cognitive functioning, ascertained later in development. Independent of model specification, there was little evidence for novel LRA-related genetic influences emerging during mid-childhood and adolescence. Thus, despite increases in trait heritability from early childhood to adolescence, developmental variation in language and literacy skills may not fully adhere to a developmental paradigm that exclusively predicts genetic innovation during the transition from early to middle childhood^10,13^.

Instead, the identification of amplification processes is consistent with twin research reporting moderate genetic correlations between latent factors for early language (including expressive vocabulary and syntax skills between 2-4 years of age) and both mid-childhood and/or adolescent latent language^10^ and reading^9^. For example, latent factors for early language explained ∼12% of the phenotypic variation in a latent factor for mid-childhood reading^9^ using individual pathway models. Based on bivariate heritability patterns between latent factors, accounting only for about a third of phenotypic correlations^9,10^, findings have been interpreted as evidence for novel genetic influences emerging during mid-childhood^10^. In the present study, early vocabulary-related genetic factors could explain up to 45.1% phenotypic variation in subsequent LRAs, especially for literacy and verbal cognition, accounting for the majority of SNP-h^2^ (≤95%). Bivariate heritability estimations confirmed these findings. Similar amplification patterns were observed between early vocabulary and PIQ, although the evidence for bivariate heritability with PIQ was less strong. This suggests that genetic variance between early vocabulary and subsequent verbal cognition and literacy, but also non-verbal cognition, is shared, showing developmental genetic stability. However, the striking similarity among structural models for many literacy skills may partially reflect their complex phenotypic interrelatedness.

The largest amplification of genetic variation contributing to later literacy and cognition was identified for a small proportion of genetic influences that is unique to receptive and independent of expressive vocabulary at 38 months of age. Consistently, bivariate heritabilities with early receptive vocabulary accounted for 70-100% of the phenotypic covariance with later reading and cognition skills, although the 95% confidence intervals are wide. In contrast, genetic influences for expressive vocabulary did not substantially contribute to the total genetic variance of later LRAs (based on their factorial co-heritability) except for VIQ scores. Analysing reverse GSEM (where the order of early vocabulary scores is reversed) confirmed these patterns. It is noteworthy that in these reverse GSEM models we also identified evidence for the amplification of a genetic factor that is unique to expressive vocabulary (i.e. independent of receptive vocabulary). However, there was little evidence for a substantial contribution to LRA SNP-h^2^. In addition, the identified bivariate heritability patterns remained unchanged. Thus, our results suggest that genetic variance between early vocabulary and subsequent literacy and cognitive skills is not only shared, but that genetic links are dominated by early receptive vocabulary, suggesting specificity, and thus only partially adhere to the concept of ‘generalist genes’^16^. Genetic links with expressive vocabulary still exist, albeit to a lower extent.

The observed differences in genetic overlap with LRAs may reflect differential mechanisms that link receptive and expressive vocabulary-related genetic factors to later reading and cognitive skills. For example, receptive vocabulary may be more strongly related to pre-reading skills, such as phonological awareness and orthographic knowledge, while expressive vocabulary has been previously identified as predictive of word identification^38^. Furthermore, a delay in both expressive and receptive vocabulary is much more likely to lead to problems with later literacy compared to delays in expressive vocabulary alone^39^.

The methodology applied in this study does not allow us to infer specific biological pathways or specific genes encoded by the identified genetic factors. However, it is still possible to speculate about the biological mechanisms that may underlie the observed amplification patterns. Genes are known to have multiple biological functions (pleiotropy), and dynamic gene expression patterns over time and space have been shown for multiple brain-related gene expression modules^40^. The stability of genetic factors across development is furthermore consistent with signalling pathways and genes that contribute to synaptic function and plasticity with important biological roles throughout development^41^, though specifically designed gene-based studies are warranted to confirm such claims.

The increase in SNP-h^2^, comparing early vocabulary skills with later language, literacy and cognitive performance, as observed in this study, may not necessarily involve an increase in genetic variance over time. Instead, it may arise due to genotype-environment correlations, implying an amplification of small genetic differences as children develop, because of environment modification and selection in accordance with their genetic make-up^42^. Furthermore, environmental variance in LRAs may decrease with the start of schooling^43^ and parent-reported vocabulary measures might be associated with higher random error rates compared to direct assessments of language and literacy measures using standardised psychological instruments, affecting heritability estimations^44^.

It should be noted that our findings do not preclude the emergence of novel genetic influences during later childhood and adolescence. Parent-reported vocabulary measures in ALSPAC have sufficient power (80%) power to detect SNP-h^2^ estimates of 0.15 (Supplementary Methods). However, compared to large-scale genome-wide studies of educational attainment^45^ or direct assessments of language and literacy measures their predictive power is low. Indeed, anthropometric measures that are more reliably assessed, such as head circumference (known to be genetically correlated with cognitive functioning), show evidence for both amplification and innovation processes from infancy to later adolescence^46^. A further limitation of the current study is that the CDI Toddler version was developed to assess vocabulary in children up to 30 months^20^, whereas ALSPAC children were assessed at 38 months of age, potentially leading to ceiling effects.

The strength of this work lies in the identification of amplification processes exploiting a temporal sequence of events, suggesting that the developmental origins of later complex cognitive and literacy processes lie in early childhood. Our findings suggest that cheaply and easily administered parent-reported CDI questionnaires, which are widely used to assess children’s early language^47^, can be useful instruments to capture common genetic influences affecting individual differences in LRAs many years later in life. Moreover, when applied to large numbers of participants (hundreds of thousands), these parent-reports could become sensitive genetic prediction tools. However, there is a need to improve their predictive power, although moderate to strong correlations between parental judgements of a child’s vocabulary and direct assessments of a child’s vocabulary suggest instrument validity^48,49^.

In summary, we show that the amplification of a small proportion of genetic influences that uniquely capture early receptive vocabulary play a major role during later cognitive and literacy development. This suggests genetic stability, with developmental origins of complex cognitive and literacy skills arising early in childhood.

## Data availability

The data used is available through a fully searchable data dictionary (http://www.bris.ac.uk/alspac/researchers/data-access/data-dictionary/). Access to ALSPAC data can be obtained as described within the ALSPAC data access policy (http://www.bristol.ac.uk/alspac/researchers/access/).

## Code availability

All analyses were performed using freely accessible software. Requests for scripts or other analysis details can be sent via email to the corresponding authors at.

## Acknowledgements

We are extremely grateful to all the families who took part in this study, the midwives for their help in recruiting them, and the whole ALSPAC team, which includes interviewers, computer and laboratory technicians, clerical workers, research scientists, volunteers, managers, receptionists and nurses. The UK Medical Research Council and Wellcome (Grant ref: 102215/2/13/2) and the University of Bristol provide core suport for ALSPAC. GWAS data was generated by Sample Logistics and Genotyping Facilities at Wellcome Sanger Institue and LabCorp (Laboratory Corporation of America) using support from 23andMe. A comprehensive list of grants funding is available on the ALSPAC website (http://www.bristol.ac.uk/alspac/external/documents/grant-acknowledgements.pdf). EV, BSTP and SEF are supported by the Max Planck Society. BSTP is also supported by the Simons Foundation (Award ID: 514787). This publication is the work of the authors and EV and BSTP will serve as guarantors for the contents of this paper.

## Author contributions

BSTP developed the study concept and EV contributed to the study design. EV performed the data analysis and interpretation under the supervision of BSTP. EV and BSTP drafted the manuscript and CYS, SEF and PSD provided critical revisions. All authors approved the final version of the manuscript for submission.

