## Supplementary Information for "The amplification of genetic factors for early vocabulary during children’s language and literacy development"

#### Supplementary Methods

ALSPAC description

14,541 pregnant women resident in Avon, UK with expected dates of delivery 1st April 1991 to 31st December 1992 were recruited by ALSPAC. Initially, 14,541 pregnancies were enrolled for which the mother enrolled in the ALSPAC study and had either returned at least one questionnaire or attended a "Children in Focus" clinic by 19/07/99. This comprised a total of 14,676 fetuses, resulting in 14,062 live births and 13,988 children who were alive at the age of one year.

An attempt was made to bolster the initial sample, when the oldest children were approximately 7 years of age, with eligible cases who had failed to join the study originally. Consequently, there are data available for more than 14,541 pregnancies (see above) when considering variables collected from the age of seven onwards (and potentially abstracted from obstetric notes).

There are 913 pregnancies not in the initial sample (known as Phase I enrolment) that are currently represented on the built files and reflecting enrolment status at the age of 24. Of these new pregnancies, 452, 262 and 195 were recruited during Phases II, III and IV respectively. As a result, an additional 913 children are enrolled. The cohort profile paper[1] describes the phases of enrolment in more detail.

Thus, the total sample size for analyses using any data collected after the age of seven is 15,454 pregnancies, resulting in 15,589 fetuses. Of this total sample 14,901 children were alive at the age of one year.

The Children in Focus (CiF) group, a 10% sample of the ALSPAC cohort, attended clinics at the University of Bristol at various time intervals between 4 to 61 months of age. The CiF group were randomly chosen from the last 6 months of ALSPAC births (1,432 families attended at least one clinic).

Mothers that had moved out of the area, were lost to follow-up, or those partaking in another study of infant development in Avon were excluded.

Please note that the study website contains details of all the data that is available through a fully searchable data dictionary and variable search tool (http://www.bristol.ac.uk/alspac/researchers/our-data).

Genetic quality control

ALSPAC participants were genotyped using the Illumina HumanHap550 quad chip genotyping platforms, and genotypes were called using the Illumina GenomeStudio software. Standard genomic quality control[2] was performed at both the SNP and individual level using PLINK (v1.07)[3]. Individuals with a gender mismatch (comparing reported gender with genetic gender), a large number of missing SNPs (>3%), non-European ancestry and a genetic relationship >0.05 were excluded from the analyses. SNPs that had a low call rate (<99%), were rare (<1%) or deviated from Hardy-Weinberg equilibrium ($P<5x10^{-7}$) were also excluded from the analysis. After quality control, 8,226 children and 465,740 SNPs remained.

Power analyses

For this study, we selected early vocabulary measures (15-38 months) that had a sample size >6,000. This corresponds to at least 80% power to detect a SNP-$h^2$ of 0.15 ($P$>0.05)[4]. For LRAs assessed later in life (7-13 years), sample sizes were slightly lower and measures that had at least 80% power to detect a SNP-$h^2$ of >0.20 ($P$>0.05)[4] were selected, corresponding to a sample size of ~4,000.

ALSPAC measures from mid-childhood to adolescence

*Reading accuracy and comprehension age 7 (WORD)*

To assess decoding and word reading both pictures and words were used. A series of four pictures, with for each picture four short, simple words underneath it, was shown to the child. Then, the child was asked to indicate, by pointing, the word that had the same beginning or ending sound as the picture. Next, a series of three pictures, again each with four words beneath starting with the same letter as the picture were shown to the child. The child was asked to point to the word underneath each picture that correctly named the picture. Finally, basic reading was assessed using the basic reading subtest of the Wechsler Objective Reading Dimensions (WORD)[5]. In summary, the child was asked to read aloud a series of 48 unconnected words which increased in difficulty. If the child made six consecutive errors, the task was stopped. The reading accuracy and comprehension score was computed as the sum of the number of items that the child read/responded to correctly.

*Reading accuracy age 9 (NBO)*

The child was asked to read aloud ten real words, selected from a larger selection of words as described by Nunes, Bryant and Olsen (NBO)[6]. The test-retest reliability of this assessment for word reading was 0.80 and a 0.85 correlation with the Schonell Word Reading Task[7] was observed. A score indicating reading accuracy was computed as the sum of the number of items that the child read correctly.

*Reading speed and reading accuracy age 9 (NARA II)*

The child was asked to read a passage from a booklet, following the revised Neale Analysis of Reading Ability (NARA II)[8]. The tester recorded both the time it took the child to read the passage, and also noted any errors made by the child. All scores were standardised by age.

*Reading speed age 13 (TOWRE)*

The child had 45 seconds to read as many words as possible from the Test of Word Reading Efficiency (TOWRE)[9] to assess sight word efficiency. The tester marked words that a child skipped, or got wrong. A reading speed score was computed as the sum of the number of correct words a child finished on.

*Non-word reading accuracy age 9 (NBO)*

The child was asked to read aloud ten non-words, selected from a larger selection of non-words taken from research conducted by Nunes and colleagues[6]. The test -retest reliability of the non-word reading task was 0.73 and correlation with the Schonell Word Reading Task[7] of 0.73 was observed. The tester emphasised to the child that the words ware made-up, and asked the child to read all the non-words in the way that they thought they should be read. A non-word reading accuracy score was computed as the sum of the number of items the child read correctly.

*Non-word reading speed age 13 (TOWRE)*

The child was asked to read as many non-words as possible within 45 seconds. Word lists were derived from the non-word part of the Test of Word Reading Efficiency (TOWRE)[9] to assess decoding efficiency. The tester marked words that a child skipped, or got wrong. A non-word reading speed score is computed as the sum of the number of correct non-words a child finished on.

*Spelling accuracy age 7 (NB)*

The child was asked to write down the spelling for a series of 15 words, chosen specifically for this age group after piloting on several hundred children (Nunes and Bryant, ALSPAC-specific measure). The words were of different frequencies, included regular and irregular words, and increased in difficulty. For each word, the tester first read only the word to the child, then a specific sentence incorporating the word, and finally alone again. A spelling accuracy score is computed as the number of words spelt correctly.

*Spelling accuracy age 9 (NB)*

Spelling accuracy at age 9 was assessed in a similar manner to that at age 7 (see above). However, the series of 15 words that a child was asked to spell were adjusted to match the age group of 9. A spelling accuracy score is computed as the number of words spelt correctly.

*Phonemic awareness age 7 (AAT)*

The task consisted of two practice and 40 test items of increasing difficulty, according to the Auditory Analysis Test (AAT)[10]. For each item, the child was asked to first repeat the word, and then produce it again but without part of the word (a phoneme or a number of phonemes). There were seven omission categories: 1) omission of a first syllable, 2) omission of a medial syllable, 3) omission of a final syllable, 4) omission of the initial consonant of a one syllable word, 5) omission of the final consonant of a one syllable word, 6) omission of the first consonant of a medial consonant, and 7) omission of the consonant blend of a medial consonant. Words from similar categories were not clustered. A phonemic awareness score is computed as the sum of correct responses.

*Listening comprehension age 8 (WOLD)*

A picture was shown to the child and the tester read aloud a paragraph about the picture, following a subset of the Wechsler Objective Language Dimensions (WOLD)[11] test. Next, the child was asked to answers ten questions on what they heard. A listening comprehension score is calculated as the sum of the items that the child got correct.

*Non-word repetition age 8 (CNRep)*

The child was asked to listen to a series of 12 nonsense words, according to an adaptation of the Children's Test of Nonword Repetition (CNRep)[12]. The 12 nonsense words consisted of four nonsense words of three syllables, four nonsense words of four syllables, and four nonsense words of five syllables. All nonsense words were conforming to English rules for sound combinations. For each words, the child was asked to repeat the word after listening to it. The repetition attempt was scored as correct if there was no phonological deviation from the target form. A non-word repetition score was computed as the sum of the number of correct non-words.

*Verbal intelligence age 8 (WISC-III)*

A short form of the Wechsler Intelligence Scale for Children (WISC-III)[13], including alternate items for all subtests, with the exception of the coding subtest was administered. The WISC-III comprises ten subtests five of which are verbal subtests: information, similarities, arithmetic, vocabulary, comprehension, and can be used to construct a verbal intelligence score. Based on the items used in the alternate item form of the WISC-III raw scores were calculated and the total age-scaled scores for the verbal scale were calculated using the look-up tables provided in the WISC-III manual. All scores were pro-rated.

*Performance intelligence age 8 (WISC-III)*

A short form of the Wechsler Intelligence Scale for Children (WISC-III)[13], including alternate items for all subtests, with the exception of the coding subtest was administered. The WISC-III contains five performance subtests: picture completion, coding, picture arrangement, block design and object assembly. Based on the items used in the alternate item form of the WISC-III raw scores were calculated and the total age-scaled scores for the performance scale were calculated using the look-up tables provided in the WISC-III manual. All scores were pro-rated.

Structural equation modelling

A Cholesky decomposition can be described as follows[14]: for a multivariate trait P with phenotypic measurements t, a latent genetic factor ($A_1$) influences the first measure $P_1$, but may also explain variance in the remaining measures ($P_2$,...,$P_t$). Additionally, a second latent genetic factor ($A_2$) influences the second measure ($P_2$) and may explain variance, not yet captured by $A_1$, in all other measures ($P_3$,...,$P_t$). The final measure ($P_t$) is influenced by latent genetic factors ($A_1$,...,$A_{t-1}$), but also a genetic factor $A_t$. This latter genetic factor does not explain variance within any of the previous measures ($P_1$,...,$P_{t-1}$)[15]. Genetic factor loadings (path coefficients) were annotated with *a.* Here, the first number indicates the direction of the effect (the variable to which the arrow points) and the second the origin of the effect[15].

The expected phenotypic covariance matrix $\mathbf{\Sigma}$ for Z-standardised traits, based on the factor model is

$$\mathbf{\Sigma} = \mathbf{\Lambda \Phi \Lambda' + \Gamma \Theta \Gamma'} \tag{1}$$

with a lower triangular matrix of genetic factor loadings $\mathbf{\Lambda}$, a diagonal matrix of latent genetic factor variances $\mathbf{\Phi}$ (standardised to unit variance), such that $\mathbf{\Phi}$ is an identity matrix ***I*** [16]. The residual variance can be decomposed into latent residual factors, with a lower triangular matrix of residual factor loadings $\mathbf{\Gamma}$ and a diagonal matrix of latent residual factor variances $\mathbf{\Theta}$ (standardised to unit variance), such that $\mathbf{\Theta}$ is an identity matrix ***I***. For example, a trivariate model consisting of three measures ($P_1$ , $P_2$ and $P_3$), assuming three genetic factors ($A_1$, $A_2$ and $A_3$) and three residual factors ($E_1$, $E_2$ and $E_3$). The expected phenotypic covariance matrix for this trivariate model can be expressed as follows:

$$\mathbf{\Sigma} = \begin{bmatrix} \sigma_{p1}^2 & \sigma_{p12} & \sigma_{p13} \\ \sigma_{p12} & \sigma_{p2}^2 & \sigma_{p23} \\ \sigma_{p13} & \sigma_{p23} & \sigma_{p3}^2 \end{bmatrix} \tag{2}$$

with the relevant matrices

$$\mathbf{\Lambda} = \begin{bmatrix} a_{11} & 0 & 0 \\ a_{21} & a_{22} & 0 \\ a_{31} & a_{32} & a_{33} \end{bmatrix}, \ \mathbf{\Phi} = \begin{bmatrix} 1 & 0 & 0 \\ 0 & 1 & 0 \\ 0 & 0 & 1 \end{bmatrix}, \ \mathbf{\Gamma} = \begin{bmatrix} e_{11} & 0 & 0 \\ e_{21} & e_{22} & 0 \\ e_{31} & e_{32} & e_{33} \end{bmatrix}, \ \mathbf{\Theta} = \begin{bmatrix} 1 & 0 & 0 \\ 0 & 1 & 0 \\ 0 & 0 & 1 \end{bmatrix} \tag{3}$$

with phenotypic variances $\sigma^2_{p1}, \sigma^2_{p2}$ and $\sigma^2_{p3}$ and phenotypic covariances $\sigma_{p12}$, $\sigma_{p13}$ and $\sigma_{p23}$.

The trivariate AE Cholesky decomposition of three standardised measures (see above), can be visualised using a path diagram (Supplementary Figure 3). The expected phenotypic variances and covariances can be expressed as follows:

$$\sigma^2_{p1} = a^2_{11} + e^2_{11} = 1 \tag{4}$$

$$\sigma^2_{p2} = (a^2_{21} + a^2_{22}) + (e^2_{21} + e^2_{22}) = 1 \tag{5}$$

$$\sigma^2_{p3} = (a^2_{31} + a^2_{32} + a^2_{33}) + (e^2_{31} + e^2_{32} + e^2_{33}) = 1 \tag{6}$$

$$\sigma_{p12} = a_{11}a_{21} + e_{11}e_{21} \tag{7}$$

$$\sigma_{p13} = a_{11}a_{31} + e_{11}e_{31} \tag{8}$$

$$\sigma_{p23} = a_{31}a_{21} + a_{32}a_{22} + e_{31}e_{21} + e_{32}e_{22} \tag{9}$$

The variance of the latent genetic and residual factors has been standardised to unit variance and is not shown.

Genetic correlation estimates between phenotypes, measuring the extent to which two phenotypes 1 and 2 share genetic factors (ranging from -1 to 1), can be derived using estimated genetic variances and covariances[17] according to:

$$r_g = \frac{\sigma_{g12}}{\sqrt{\sigma_{g1}^2 \sigma_{g2}^2}} \tag{10}$$

with genetic covariance $\sigma_{g12}$ between phenotypes 1 and 2 and the genetic variances $\sigma_{g1}^2$ and $\sigma_{g2}^2$.

Factorial co-heritability

Factorial co-heritability estimates the proportion of total genetic variance observed for a trait that is accounted for by a specific genetic factor, and was estimated using the gsem package (R:gsem library, version 0.1.5). Factorial co-heritability was calculated based on standardised path coefficients, corresponding SEs were derived using the Delta method, and *P*-values approximated with a Wald test.

Bivariate heritability

Bivariate heritability[18] details the proportion of phenotypic covariance between two traits that is accounted for by the genetic covariance, and was estimated using the gsem package (R:gsem library, version 0.1.5). The genetic covariance was estimated based on unstandardised path coefficients and the phenotypic covariance on rank-transformed measures. SEs were approximated by the SE of the genetic covariance divided by the phenotypic covariance (as the SE of the phenotypic covariance is small) and *P*-values based on a Wald-test, assuming normality. Reported bivariate heritability estimates are based on forward GSEM models, and reverse GSEM models provided nearly identical results.

Websites

GCTA: https://cnsgenomics.com/software/gcta/

GSEM: https://gitlab.gwdg.de/beate.stpourcain/gsem

Metafor: http://www.metafor-project.org/doku.php

matSpD: https://gump.qimr.edu.au/general/daleN/matSpD/

#### Supplementary Tables

##### Supplementary Table 1: SNP-heritability estimates

| Measure | N | GCTA-$h^2$ (SE) | GSEM-$h^2$ (SE) |
|---|---|---|---|
| Expressive voc 38m (CDI) | 6,092 | 0.18(0.06) | 0.18(0.06)* |
| Receptive voc 38m (CDI) | 6,092 | 0.12(0.06) | 0.13(0.04)* |
| Reading a/c 7 (WORD) | 5,723 | 0.42(0.06) | 0.41(0.06) |
| Reading a 9 (NBO) | 5,574 | 0.46(0.06) | 0.46(0.06) |
| Reading a 9 (NARA II) | 5,048 | 0.50(0.07) | 0.49(0.07) |
| Reading s 9 (NARA II) | 5,037 | 0.45(0.07) | 0.43(0.07) |
| Reading s 13 (TOWRE) | 4,131 | 0.40(0.09) | 0.41(0.09) |
| NW reading a 9 (NBO) | 5,569 | 0.32(0.06) | 0.33(0.06) |
| NW reading s 13 (TOWRE) | 4,121 | 0.38(0.06) | 0.38(0.09) |
| Spelling a 7 (NB) | 5,637 | 0.32(0.06) | 0.33(0.06) |
| Spelling a 9 (NB) | 5,564 | 0.38(0.06) | 0.38(0.07) |
| PhonAware 7 (AAT) | 5,749 | 0.39(0.06) | 0.38(0.06) |
| Listening c 8 (WOLD) | 5,324 | 0.32(0.07) | 0.30(0.07) |
| NW repetition (CNRep) | 5,315 | 0.32(0.07) | 0.31(0.06) |
| VIQ 8 (WISC-III) | 5,305 | 0.54(0.07) | 0.54(0.06) |
| PIQ 8 (WISC-III) | 5,296 | 0.26(0.07) | 0.26(0.06) |

* GSEM-$h^2$ estimates as observed in the GSEM model with verbal intelligence quotient at 8 years.

Abbreviations: a, accuracy; AAT, Auditory Analysis Test; c, comprehension; CDI, Communicative Development Inventory; CNRep, Children's Test of Nonword Repetition; GCTA, genome-wide complex trait analysis; GSEM, Genetic-relationship-matrix Structural Equation modelling; $h^2$, heritability; m, months; NARA II, The Neale Analysis of Reading Ability- Second Revised British Edition; NB, ALSPAC-specific assessment developed by Nunes and Bryant; NBO, ALSPAC-specific assessment developed by Nunes, Bryant and Olson; NW, nonword; PhonAware, phonemic awareness; PIQ; performance intelligence quotient; s, speed; TOWRE, Test Of Word Reading Efficiency; VIQ, verbal intelligence quotient; voc, vocabulary; WISC-III, Wechsler Intelligence Scale for Children III; WOLD, Wechsler Objective Language Dimensions; WORD, Wechsler Objective Reading Dimension

SNP-heritability estimates were estimated based on rank-transformed scores, directly genotyped SNPs and individuals with a genetic relationship of <0.05, using Restricted Maximum Likelihood (REML) analyses as implemented in genome-wide complex trait analysis (GCTA) software. SNP-heritability estimates based on forward Genetic-relationship-matrix Structural Equation modelling (GSEM) were extracted for comparison.

Supplementary Table 2: Meta-analysis domains of mid-childhood to adolescence language- and literacy-related abilities

| LRA | Meta-analysis domains | | |
|---|---|---|---|
| | All LRAs | Reading | Spelling |
| Reading a/c 7 (WORD) | ✓ | ✓ | × |
| Reading a 9 (NBO) | ✓ | ✓ | × |
| Reading a 9 (NARA II) | ✓ | ✓ | × |
| Reading s 9 (NARA II) | ✓ | ✓ | × |
| Reading s 13 (TOWRE) | ✓ | ✓ | × |
| NW reading a 9 (NBO) | ✓ | ✓ | × |
| NW reading s 13 (TOWRE) | ✓ | ✓ | × |
| Spelling a 7 (NB) | ✓ | × | ✓ |
| Spelling a 9 (NB) | ✓ | × | ✓ |
| PhonAware 7 (AAT) | ✓ | × | × |
| Listening c 8 (WOLD) | ✓ | × | × |
| NW repetition (CNRep) | ✓ | × | × |
| VIQ 8 (WISC-III) | ✓ | × | × |

Abbreviations: a, accuracy; AAT, Auditory Analysis Test; c, comprehension; CNRep, Children's Test of Nonword Repetition; LRAs, language- and literacy-related abilities; NARA II, The Neale Analysis of Reading Ability- Second Revised British Edition; NB, ALSPAC-specific assessment developed by Nunes and Bryant; NBO, ALSPAC-specific assessment developed by Nunes, Bryant and Olson; NW, nonword; PhonAware, phonemic awareness; s, speed; TOWRE, Test Of Word Reading Efficiency; VIQ, verbal intelligence quotient; WISC-III, Wechsler Intelligence Scale for Children III; WOLD, Wechsler Objective Language Dimensions; WORD, Wechsler Objective Reading Dimension

Absolute path coefficients estimated using genetic-relationship-matrix structural equation (GSEM) models were meta-analysed, accounting for trait-interrelationships. Meta-analysis were carried out across all LRAs, and for reading-related abilities as well as spelling-related abilities. ✓ indicates that a specific trait was included in a meta-analysis, × indicates that a specific trait was not included.

Supplementary Table 3: Factorial co-heritabilities

| Measure | Forward GSEM | | | | Reverse GSEM | | | |
|---|---|---|---|---|---|---|---|---|
| | Expressive vocabulary 38 months* | | Receptive vocabulary 38 months** | | Expressive vocabulary 38 months*** | | Receptive vocabulary 38 months **** | |
| | Factorial co-heritability (SE) | *P* | Factorial co-heritability (SE) | *P* | Factorial co-heritability (SE) | *P* | Factorial co-heritability (SE) | *P* |
| Reading a/c 7 (WORD) | 0.06(0.07) | 0.41 | 0.94(0.08) | $<1x10^{-10}$ | 0.43(0.23) | 0.06 | 0.57(0.23) | 0.01 |
| Reading a 9 (NBO) | 0.10(0.10) | 0.29 | 0.33(0.45) | 0.47 | 0.10(0.30) | 0.73 | 0.33(0.24) | 0.17 |
| Reading a 9 (NARA II) | 0.15(0.12) | 0.19 | 0.85(0.12) | $<1x10^{-10}$ | 0.29(0.23) | 0.22 | 0.71(0.23) | 0.002 |
| Reading s 9 (NARA II) | 0.09(0.09) | 0.36 | 0.91(0.09) | $<1x10^{-10}$ | 0.40(0.24) | 0.10 | 0.60(0.23) | 0.01 |
| Reading s 13 (TOWRE) | 0.09(0.11) | 0.43 | 0.91(0.38) | 0.01 | 0.42(0.40) | 0.29 | 0.57(0.30) | 0.05 |
| NW reading a 9 (NBO) | 0.07(0.09) | 0.46 | 0.50(0.73) | 0.49 | 0.23(0.54) | 0.68 | 0.35(0.29) | 0.23 |
| NW reading s 13 (TOWRE) | 0.05(0.08) | 0.56 | 0.95(0.20) | $2x10^{-6}$ | 0.49(0.27) | 0.07 | 0.51(0.27) | 0.05 |
| Spelling a 7 (NB) | 0.09(0.10) | 0.38 | 0.90(0.37) | 0.01 | 0.45(0.27) | 0.10 | 0.55(0.27) | 0.04 |
| Spelling a 9 (NB) | 0.12(0.12) | 0.29 | 0.43(0.60) | 0.47 | 0.14(0.38) | 0.71 | 0.40(0.29) | 0.16 |
| PhonAware 7 (AAT) | 0.16(0.13) | 0.21 | 0.79(1.03) | 0.44 | 0.35(0.94) | 0.71 | 0.63(0.42) | 0.14 |
| Listening c 8 (WOLD) | 0.06(0.09) | 0.52 | 0.74(0.99) | 0.45 | 0.40(0.82) | 0.63 | 0.40(0.32) | 0.20 |
| NW repetition 8 (CNRep) | 0.20(0.16) | 0.21 | 0.54(0.81) | 0.50 | 0.17(0.53) | 0.75 | 0.58(0.40) | 0.15 |
| VIQ 8 (WISC-III) | 0.16(0.11) | 0.14 | 0.84(0.11) | $<1x10^{-10}$ | 0.27(0.20) | 0.17 | 0.73(0.20) | $2x10^{-4}$ |
| PIQ 8 (WISC-III) | 0.01(0.05) | 0.78 | 0.99(0.04) | $<1x10^{-10}$ | 0.59(0.26) | 0.02 | 0.41(0.26) | 0.11 |

Abbreviations: a, accuracy; AAT, Auditory Analysis Test; c, comprehension; CNRep, Children's Test of Nonword Repetition; GSEM, Genetic-relationship-matrix Structural Equation modelling; NARA II, The Neale Analysis of Reading Ability- Second Revised British Edition; NB, ALSPAC-specific assessment developed by Nunes and Bryant; NBO, ALSPAC-specific assessment developed by Nunes, Bryant and Olson; NW, nonword; PhonAware, phonemic awareness; PIQ; performance intelligence quotient; s, speed; TOWRE, Test Of Word Reading Efficiency; VIQ, verbal intelligence quotient; WISC-III, Wechsler Intelligence Scale for Children III; WOLD, Wechsler Objective Language Dimensions; WORD, Wechsler Objective Reading Dimension

Factorial co-heritabilities reflect the proportion of total genetic variance explained by a specific genetic factor. SEs were derived using the Delta method and *P*-values based on a Wald-test assuming normality.

* Proportion of genetic influences for expressive vocabulary including those shared with receptive vocabulary (forward GSEM model, Supplementary Figure 4) with respect to the total LRA SNP-$h^2$: $a_{31} * a_{31} / (a_{31} * a_{31} + a_{32} * a_{32} + a_{33} * a_{33})$

** Proportion of genetic influences for receptive vocabulary independent of expressive vocabulary (forward GSEM model, Supplementary Figure 4) with respect to the total LRA SNP-$h^2$: $a_{32} * a_{32} / (a_{31} * a_{31} + a_{32} * a_{32} + a_{33} * a_{33})$
*** Proportion of genetic influences for expressive vocabulary independent of receptive vocabulary (reverse GSEM model, Supplementary Figure 6) with respect to the total LRA SNP-$h^2$: $a_{32} * a_{32} / (a_{31} * a_{31} + a_{32} * a_{32} + a_{33} * a_{33})$
**** Proportion of genetic influences for receptive vocabulary including those shared with expressive vocabulary (reverse GSEM model, Supplementary Figure 6) with respect to the total LRA SNP-$h^2$: $a_{32} * a_{32} / (a_{31} * a_{31} + a_{32} * a_{32} + a_{33} * a_{33})$

The experiment-wide threshold is $P \leq 0.005$.

Supplementary Table 4: Bivariate heritability estimates

| Measure | Expressive vocabulary 38 months | | Receptive vocabulary 38 months | |
|---|---|---|---|---|
| | Bivariate heritability (SE) | *P* | Bivariate heritability (SE) | *P* |
| Reading a/c 7 (WORD) | 0.39(0.25) | 0.12 | 0.90(0.21) | $2x10^{-5}$ |
| Reading a 9 (NBO) | 0.54(0.26) | 0.04 | 0.71(0.24) | 0.003 |
| Reading a 9 (NARA II) | 0.71(0.28) | 0.01 | 1.00(0.22)* | $9x10^{-7}$ |
| Reading s 9 (NARA II) | 0.69(0.38) | 0.07 | 1.00(0.27)* | $8x10^{-10}$ |
| Reading s 13 (TOWRE) | 0.59(0.38) | 0.12 | 1.00(0.31)* | $3x10^{-4}$ |
| NW reading a 9 (NBO) | 0.40(0.28) | 0.16 | 0.71(0.28) | 0.01 |
| NW reading s 13 (TOWRE) | 0.48(0.43) | 0.26 | 1.00(0.34)* | $8x10^{-4}$ |
| Spelling a 7 (NB) | 0.59(0.35) | 0.09 | 0.95(0.26) | $3x10^{-4}$ |
| Spelling a 9 (NB) | 0.62(0.31) | 0.04 | 0.79(0.26) | 0.002 |
| PhonAware 7 (AAT) | 0.63(0.26) | 0.02 | 0.92(0.24) | $1x10^{-4}$ |
| Listening c 8 (WOLD) | 0.49(0.39) | 0.20 | 0.77(0.28) | 0.006 |
| Non-word repetition 8 (CNRep) | 0.53(0.23) | 0.02 | 0.76(0.23) | 0.001 |
| VIQ 8 (WISC-III) | 0.69(0.24) | 0.005 | 0.91(0.16) | $3x10^{-8}$ |
| PIQ 8 (WISC-III) | 0.24(0.45) | 0.59 | 0.75(0.27) | 0.006 |

Abbreviations: a, accuracy; AAT, Auditory Analysis Test; c, comprehension; CNRep, Children's Test of Nonword Repetition; NARA II, The Neale Analysis of Reading Ability- Second Revised British Edition; NB, ALSPAC-specific assessment developed by Nunes and Bryant; NBO, ALSPAC-specific assessment developed by Nunes, Bryant and Olson; NW, nonword; PhonAware, phonemic awareness; PIQ; performance intelligence quotient; s, speed; TOWRE, Test Of Word Reading Efficiency; VIQ, verbal intelligence quotient; WISC-III, Wechsler Intelligence Scale for Children III; WOLD, Wechsler Objective Language Dimensions; WORD, Wechsler Objective Reading Dimension

* Estimates were truncated at one.

Bivariate heritability estimates, reflecting the proportion of the phenotypic covariance that is accounted for by the genetic covariance. SEs were approximated by the SE of the genetic covariance divided by the phenotypic covariance (as the SE of the phenotypic covariance is small) and *P*-values are based on a Wald-test, assuming normality. Estimates are based on forward GSEM models, and reverse GSEM models provided nearly identical results (data not shown).

The experiment-wide threshold is $P \leq 0.005$.

Supplementary Figures

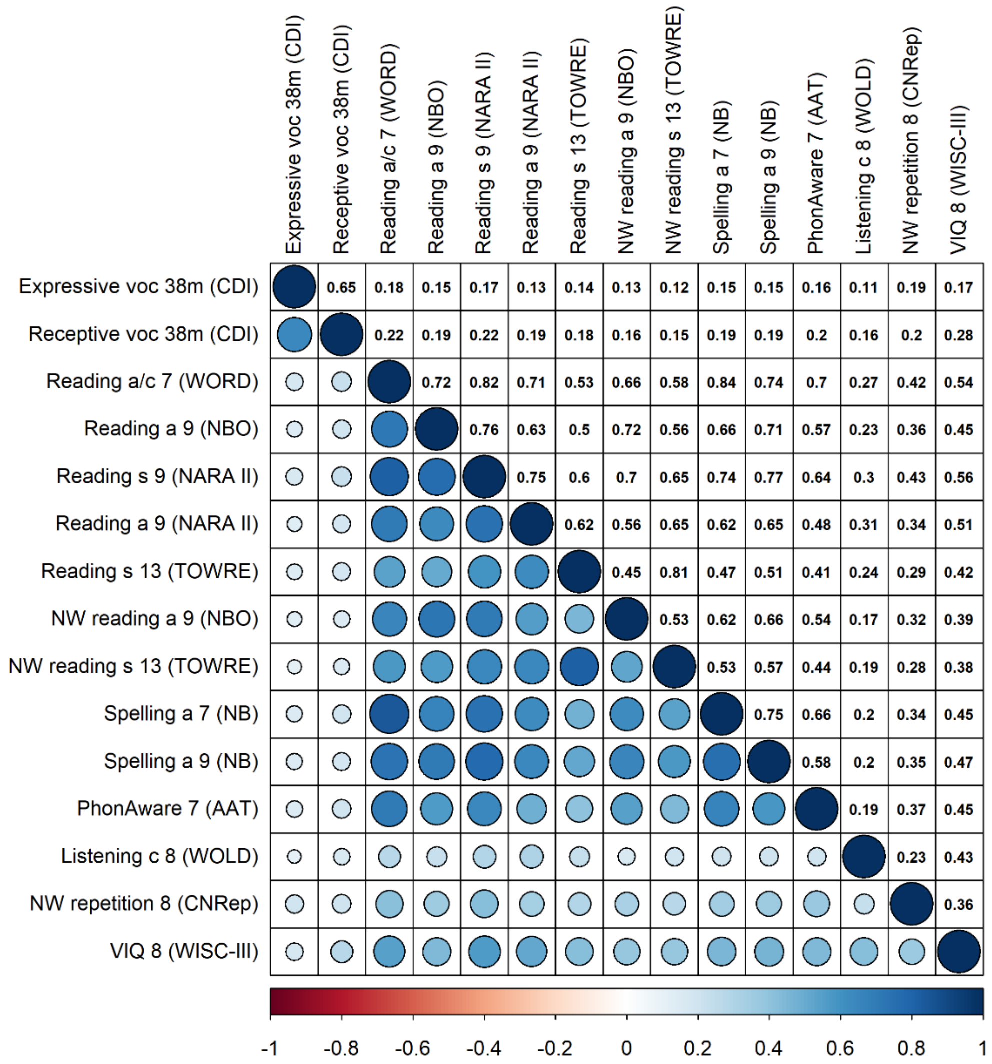

**Supplementary Figure 1: Phenotypic correlations among early vocabulary and mid-childhood to adolescence literacy- and language-related abilities (untransformed measures)**

Abbreviations: a, accuracy; AAT, Auditory Analysis Test; c, comprehension; CDI, Communicative Development Inventory; CNRep, Children's Test of Nonword Repetition; m, months; NARA II, The Neale Analysis of Reading Ability- Second Revised British Edition; NB, ALSPAC-specific assessment developed by Nunes and Bryant; NBO, ALSPAC-specific assessment developed by Nunes, Bryant and Olson; NW, nonword; PhonAware, phonemic awareness; s, speed; TOWRE, Test Of Word Reading Efficiency; VIQ, verbal intelligence quotient; voc, vocabulary; WISC-III, Wechsler Intelligence Scale for Children III; WOLD, Wechsler Objective Language Dimensions; WORD, Wechsler Objective Reading Dimension

Phenotypic correlations among untransformed measures were estimated with Spearman's rank correlation coefficients. Only correlation coefficients passing the experiment-wide significance threshold ($P \leq 0.005$) are shown.

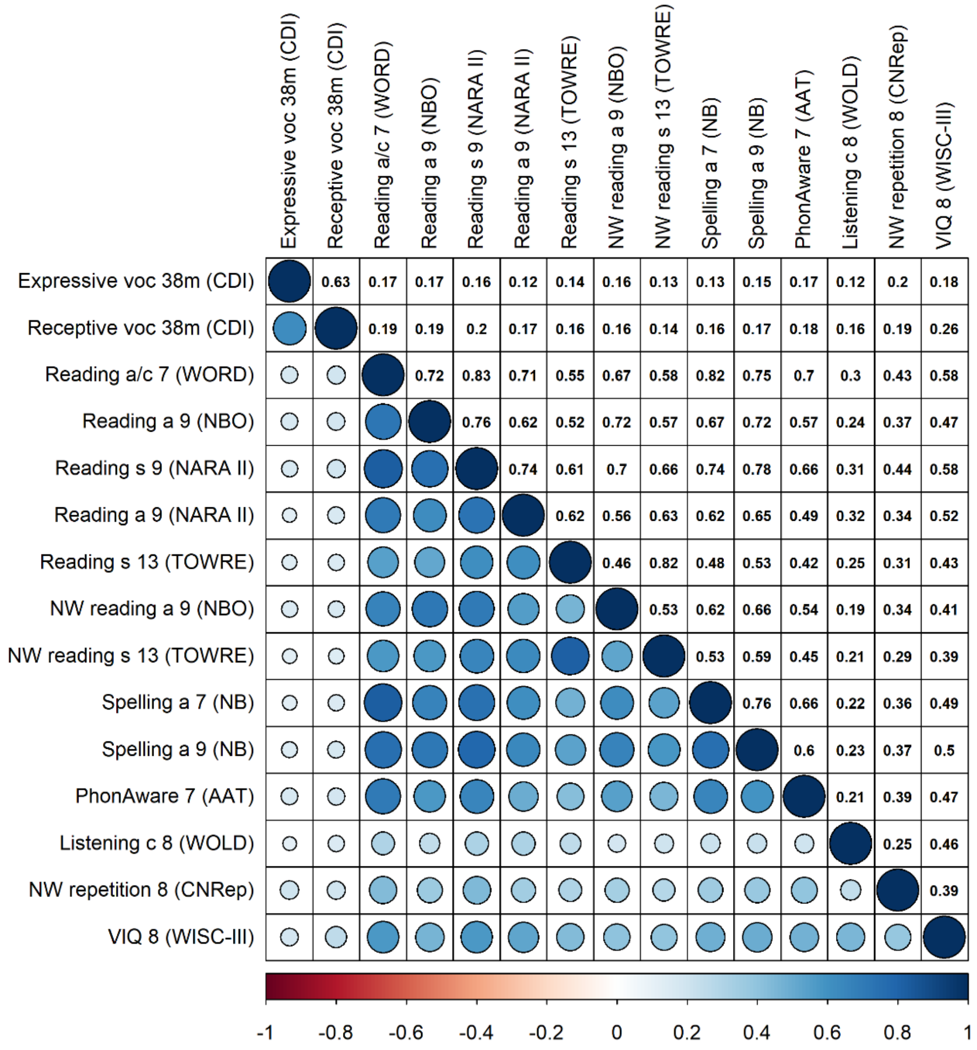

**Supplementary Figure 2: Phenotypic correlations among early vocabulary and mid-childhood to adolescence literacy- and language-related abilities (rank-transformed measures)**

Abbreviations: a, accuracy; AAT, Auditory Analysis Test; c, comprehension; CDI, Communicative Development Inventory; CNRep, Children's Test of Nonword Repetition; m, months; NARA II, The Neale Analysis of Reading Ability- Second Revised British Edition; NB, ALSPAC-specific assessment developed by Nunes and Bryant; NBO, ALSPAC-specific assessment developed by Nunes, Bryant and Olson; NW, nonword; PhonAware, phonemic awareness; s, speed; TOWRE, Test Of Word Reading Efficiency; VIQ, verbal intelligence quotient; voc, vocabulary; WISC-III, Wechsler Intelligence Scale for Children III; WOLD, Wechsler Objective Language Dimensions; WORD, Wechsler Objective Reading Dimension

Phenotypic correlations among rank-transformed measures were estimated with Pearson correlation coefficients. Only correlation coefficients passing the experiment-wide significance threshold ($P \leq 0.005$) are shown.

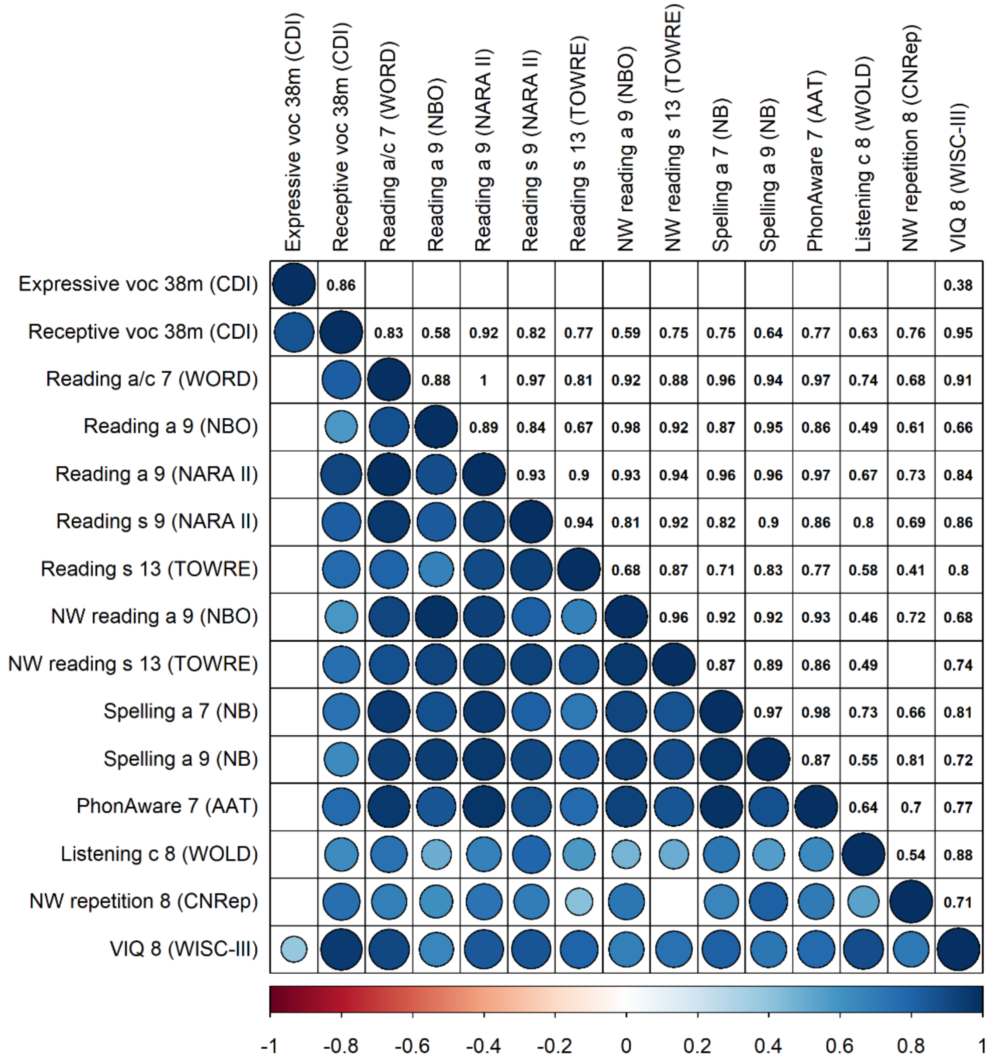

Supplementary Figure 3: Genetic correlations among early vocabulary and mid-childhood to adolescence literacy- and language-related abilities

Abbreviations: a, accuracy; AAT, Auditory Analysis Test; c, comprehension; CDI, Communicative Development Inventory; CNRep, Children's Test of Nonword Repetition; m, months; NARA II, The Neale Analysis of Reading Ability- Second Revised British Edition; NB, ALSPAC-specific assessment developed by Nunes and Bryant; NBO, ALSPAC-specific assessment developed by Nunes, Bryant and Olson; NW, nonword; PhonAware, phonemic awareness; s, speed; TOWRE, Test Of Word Reading Efficiency; VIQ, verbal intelligence quotient; voc, vocabulary; WISC-III, Wechsler Intelligence Scale for Children III; WOLD, Wechsler Objective Language Dimensions; WORD, Wechsler Objective Reading Dimension

Genetic correlations were calculated based on rank-transformed scores using Restricted Maximum Likelihood (REML) analyses as implemented in genome-wide complex trait analysis (GCTA) software, based on directly genotyped SNPs and unrelated individuals (genetic relationship of <0.05). Only genetic correlations that passed the experiment-wide significance threshold ($P \leq 0.005$) are shown.

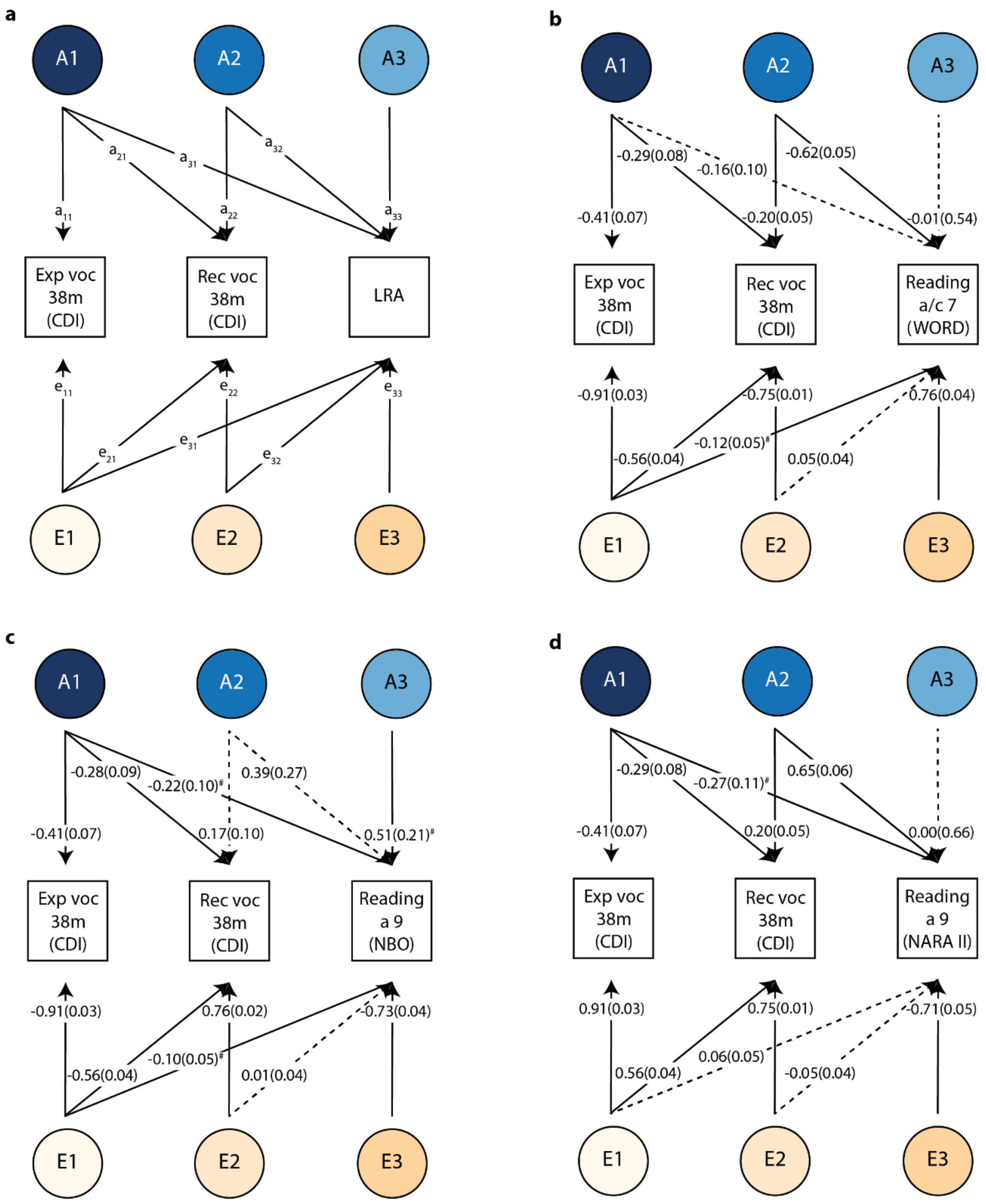
a
A1
A2
A3
a11
a21
a31
a22
a32
a33
Exp voc 38m (CDI)
Rec voc 38m (CDI)
LRA
e11
e21
e31
e22
e32
e33
E1
E2
E3
b
A1
A2
A3
-0.29(0.08)
-0.16(0.10)
-0.62(0.05)
-0.41(0.07)
-0.20(0.05)
-0.01(0.54)
Exp voc 38m (CDI)
Rec voc 38m (CDI)
Reading a/c 7 (WORD)
-0.91(0.03)
-0.75(0.01)
0.76(0.04)
-0.56(0.04)
-0.12(0.05)#
0.05(0.04)
E1
E2
E3
c
A1
A2
A3
-0.28(0.09)
-0.22(0.10)#
0.39(0.27)
-0.41(0.07)
0.17(0.10)
0.51(0.21)#
Exp voc 38m (CDI)
Rec voc 38m (CDI)
Reading a 9 (NBO)
-0.91(0.03)
0.76(0.02)
-0.73(0.04)
-0.56(0.04)
-0.10(0.05)#
0.01(0.04)
E1
E2
E3
d
A1
A2
A3
-0.29(0.08)
-0.27(0.11)#
0.65(0.06)
-0.41(0.07)
0.20(0.05)
0.00(0.66)
Exp voc 38m (CDI)
Rec voc 38m (CDI)
Reading a 9 (NARA II)
0.91(0.03)
0.75(0.01)
-0.71(0.05)
0.56(0.04)
0.06(0.05)
-0.05(0.04)
E1
E2
E3

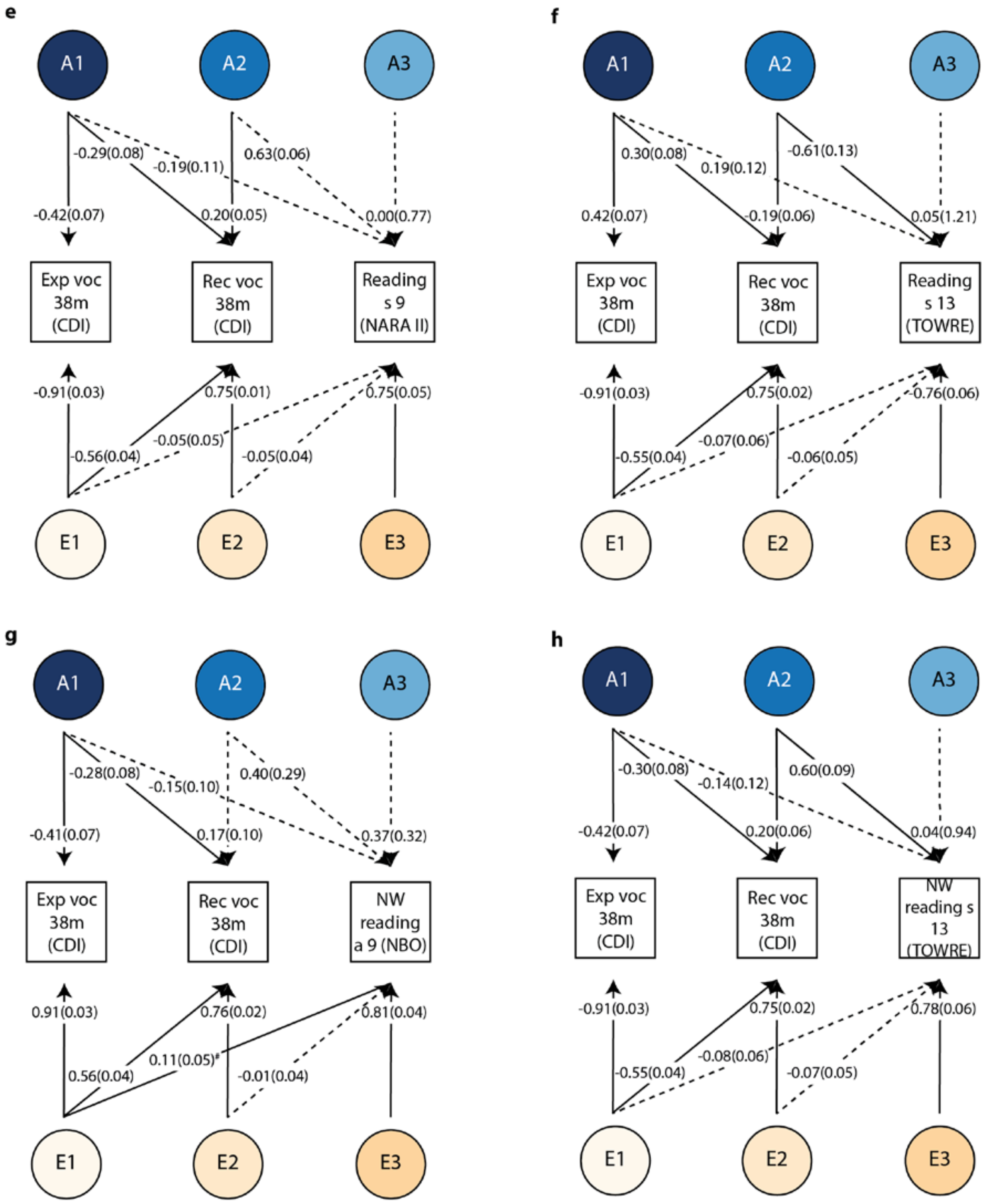
e
A1
A2
A3
-0.29(0.08)
-0.19(0.11)
0.63(0.06)
-0.42(0.07)
0.20(0.05)
0.00(0.77)
Exp voc 38m (CDI)
Rec voc 38m (CDI)
Reading s 9 (NARA II)
-0.91(0.03)
0.75(0.01)
0.75(0.05)
-0.05(0.05)
-0.56(0.04)
-0.05(0.04)
E1
E2
E3
f
A1
A2
A3
0.30(0.08)
0.19(0.12)
-0.61(0.13)
0.42(0.07)
-0.19(0.06)
0.05(1.21)
Exp voc 38m (CDI)
Rec voc 38m (CDI)
Reading s 13 (TOWRE)
-0.91(0.03)
0.75(0.02)
-0.76(0.06)
-0.07(0.06)
-0.55(0.04)
-0.06(0.05)
E1
E2
E3
g
A1
A2
A3
-0.28(0.08)
-0.15(0.10)
0.40(0.29)
-0.41(0.07)
0.17(0.10)
0.37(0.32)
Exp voc 38m (CDI)
Rec voc 38m (CDI)
NW reading a 9 (NBO)
0.91(0.03)
0.76(0.02)
0.81(0.04)
0.11(0.05)#
0.56(0.04)
-0.01(0.04)
E1
E2
E3
h
A1
A2
A3
-0.30(0.08)
-0.14(0.12)
0.60(0.09)
-0.42(0.07)
0.20(0.06)
0.04(0.94)
Exp voc 38m (CDI)
Rec voc 38m (CDI)
NW reading s 13 (TOWRE)
-0.91(0.03)
0.75(0.02)
0.78(0.06)
-0.08(0.06)
-0.55(0.04)
-0.07(0.05)
E1
E2
E3

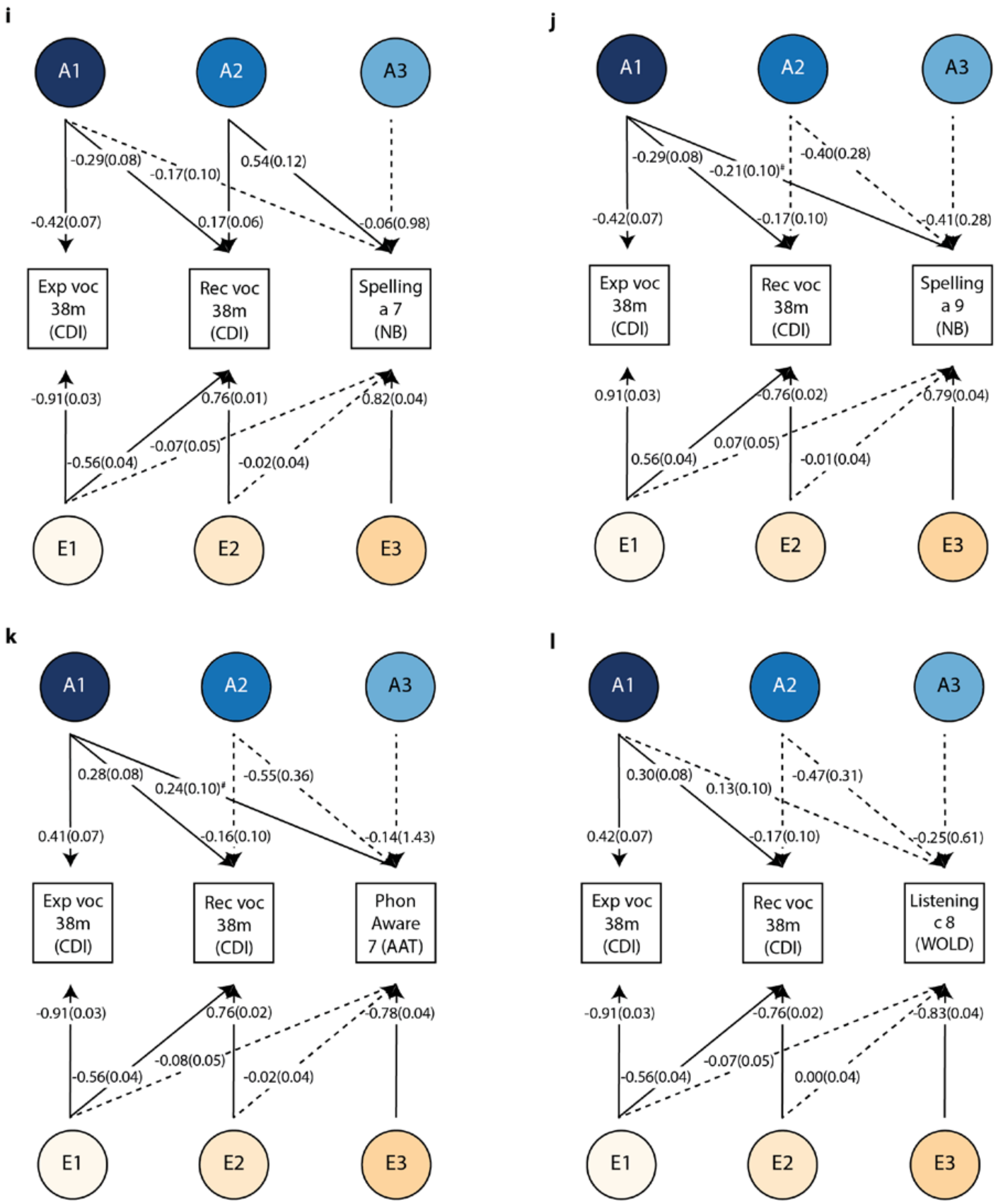
i
A1
A2
A3
-0.29(0.08)
-0.17(0.10)
0.54(0.12)
-0.42(0.07)
0.17(0.06)
-0.06(0.98)
Exp voc 38m (CDI)
Rec voc 38m (CDI)
Spelling a 7 (NB)
-0.91(0.03)
0.76(0.01)
0.82(0.04)
-0.56(0.04)
-0.07(0.05)
-0.02(0.04)
E1
E2
E3
j
A1
A2
A3
-0.29(0.08)
-0.21(0.10)#
-0.40(0.28)
-0.42(0.07)
-0.17(0.10)
-0.41(0.28)
Exp voc 38m (CDI)
Rec voc 38m (CDI)
Spelling a 9 (NB)
0.91(0.03)
-0.76(0.02)
0.79(0.04)
0.56(0.04)
0.07(0.05)
-0.01(0.04)
E1
E2
E3
k
A1
A2
A3
0.28(0.08)
0.24(0.10)#
-0.55(0.36)
0.41(0.07)
-0.16(0.10)
-0.14(1.43)
Exp voc 38m (CDI)
Rec voc 38m (CDI)
Phon Aware 7 (AAT)
-0.91(0.03)
0.76(0.02)
-0.78(0.04)
-0.56(0.04)
-0.08(0.05)
-0.02(0.04)
E1
E2
E3
l
A1
A2
A3
0.30(0.08)
0.13(0.10)
-0.47(0.31)
0.42(0.07)
-0.17(0.10)
-0.25(0.61)
Exp voc 38m (CDI)
Rec voc 38m (CDI)
Listening c 8 (WOLD)
-0.91(0.03)
-0.76(0.02)
-0.83(0.04)
-0.56(0.04)
-0.07(0.05)
0.00(0.04)
E1
E2
E3

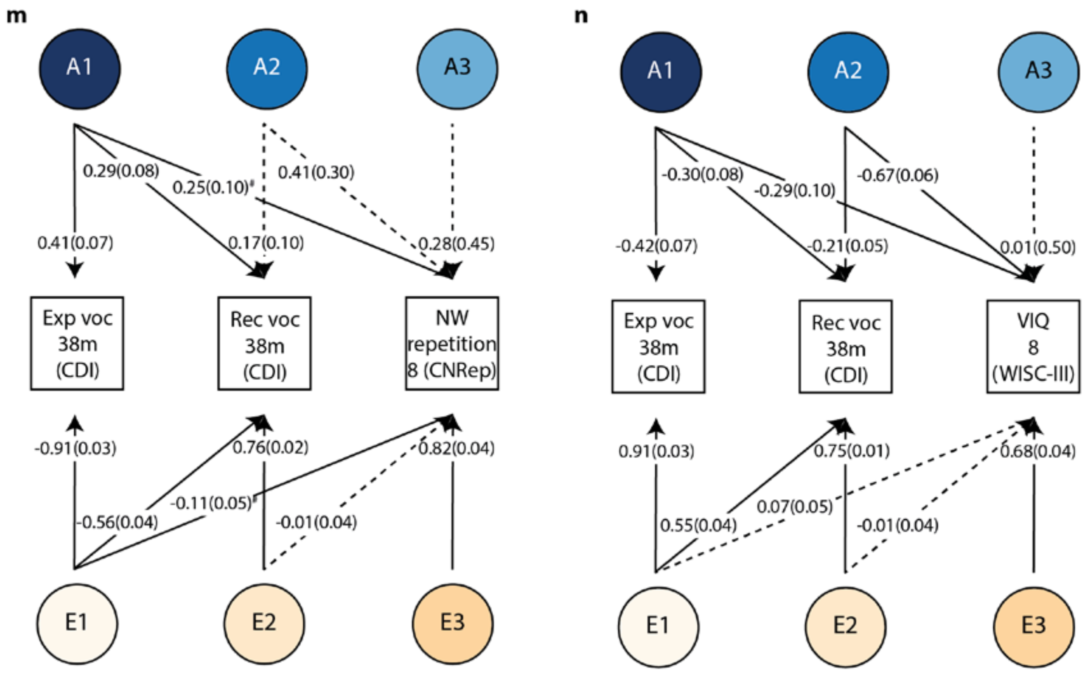

**Supplementary Figure 4: Path models of early vocabulary and mid-childhood to adolescence literacy- and language-related abilities (forward GSEM)**

Abbreviations: a, accuracy; AAT, Auditory Analysis Test; c, comprehension; CDI, Communicative Development Inventory; CNRep, Children's Test of Nonword Repetition; Exp, expressive; LRA, language- and literacy-related ability; m, months; NARA II, The Neale Analysis of Reading Ability- Second Revised British Edition; NB, ALSPAC-specific assessment developed by Nunes and Bryant; NBO, ALSPAC-specific assessment developed by Nunes, Bryant and Olson; NW, nonword; PhonAware, phonemic awareness; Rec, receptive; s, speed; TOWRE, Test Of Word Reading Efficiency; VIQ, verbal intelligence quotient; voc, vocabulary; WISC-III, Wechsler Intelligence Scale for Children III; WOLD, Wechsler Objective Language Dimensions; WORD, Wechsler Objective Reading Dimension

### Path coefficient passing nominal significance ($P \leq 0.05$), but not the experiment-wide significance threshold ($P \leq 0.005$).

Cholesky decompositions were fitted using GSEM, according to forward GSEMs and based on all available observations for children across development (N≤6,092). **(a)** Schematic path model with path coefficient labels for a Cholesky decomposition model of vocabulary at 38 months, including expressive and receptive vocabulary (in that order), and one later LRA. **(b-l)** Path models of standardised path coefficients and corresponding standard errors for 13 forward GSEM models, one for each fitted LRA. Solid lines indicate path coefficients passing a *P*-value threshold of $P \leq 0.05$, dashed lines indicate non-significant path coefficients $P > 0.05$.

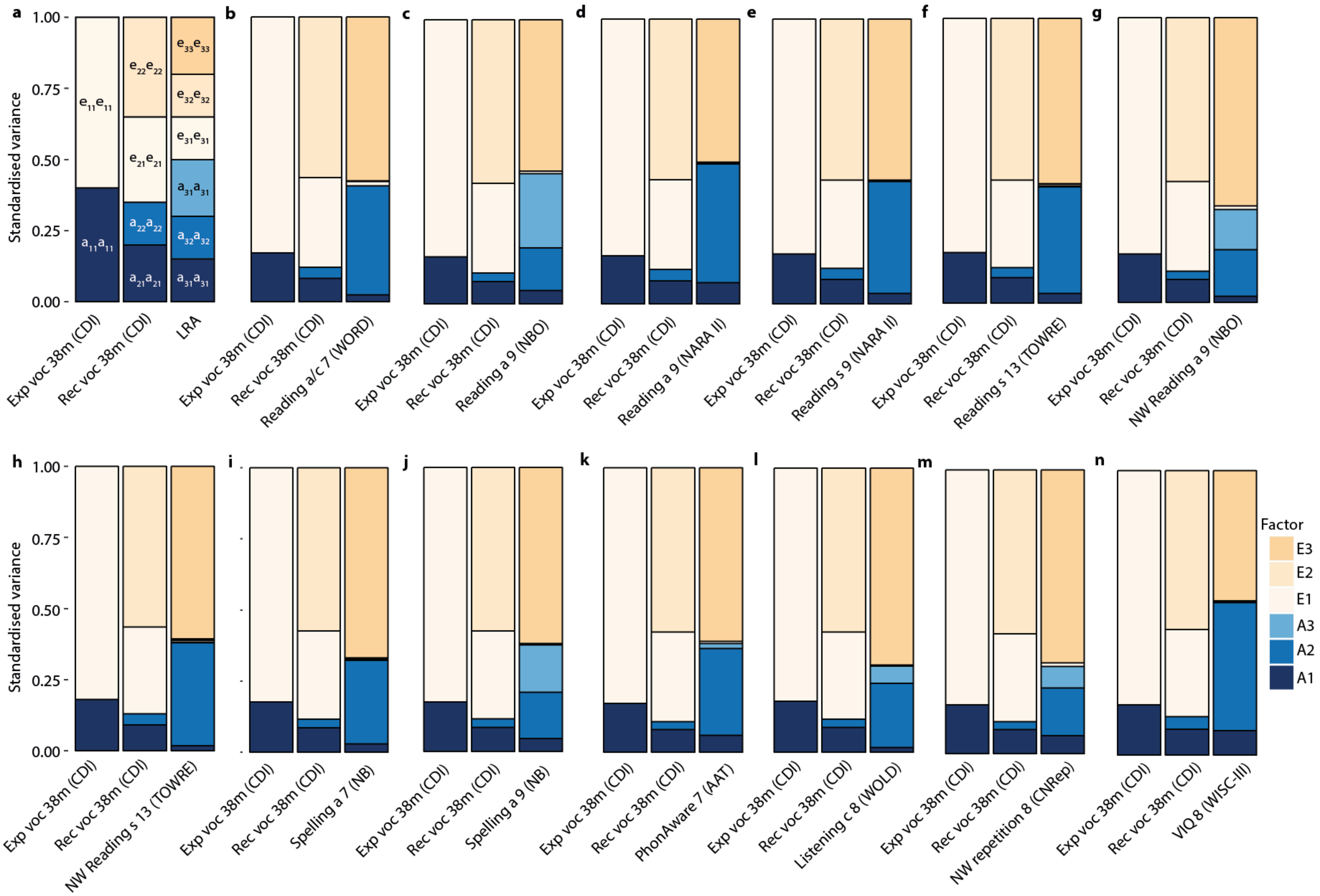
a
b
c
d
e
f
g
h
i
j
k
l
m
n
Standardised variance
1.00
0.75
0.50
0.25
0.00
e11e11
e22e22
e21e21
e33e33
e32e32
e31e31
a31a31
a32a32
a22a22
a11a11
a21a21
a31a31
Exp voc 38m (CDI)
Rec voc 38m (CDI)
LRA
Reading a/c 7 (WORD)
Reading a 9 (NBO)
Reading a 9 (NARA II)
Reading s 9 (NARA II)
Reading s 13 (TOWRE)
NW Reading a 9 (NBO)
NW Reading s 13 (TOWRE)
Spelling a 7 (NB)
Spelling a 9 (NB)
PhonAware 7 (AAT)
Listening c 8 (WOLD)
NW repetition 8 (CNRep)
VIQ 8 (WISC-III)
Factor
E3
E2
E1
A3
A2
A1

Supplementary Figure 5: Variance plots for path models of early vocabulary and mid-childhood to adolescence literacy- and language-related abilities (forward GSEM)

Abbreviations: a, accuracy; AAT, Auditory Analysis Test; c, comprehension; CDI, Communicative Development Inventory; CNRep, Children's Test of Nonword Repetition; Exp, expressive; LRA, language- and literacy-related ability; m, months; NARA II, The Neale Analysis of Reading Ability- Second Revised British Edition; NB, ALSPAC-specific assessment developed by Nunes and Bryant; NBO, ALSPAC-specific assessment developed by Nunes, Bryant and Olson; NW, nonword; PhonAware, phonemic awareness; Rec, receptive; s, speed; TOWRE, Test Of Word Reading Efficiency; VIQ, verbal intelligence quotient; voc, vocabulary; WISC-III, Wechsler Intelligence Scale for Children III; WOLD, Wechsler Objective Language Dimensions; WORD, Wechsler Objective Reading Dimension

Standardised variance explained by genetic and residual factors as derived by Cholesky decompositions using forward GSEM (Supplementary Figure 4), based on all available observations for children across development (N≤6,092). **(a)** Variance plot with path coefficient labels for a Cholesky decomposition model of vocabulary at 38 months, including expressive and receptive vocabulary (in that order), and one later LRA. **(b-l)** Standardised variance explained by genetic and residual factors as modelled in 13 forward GSEM models, one for each fitted LRA.

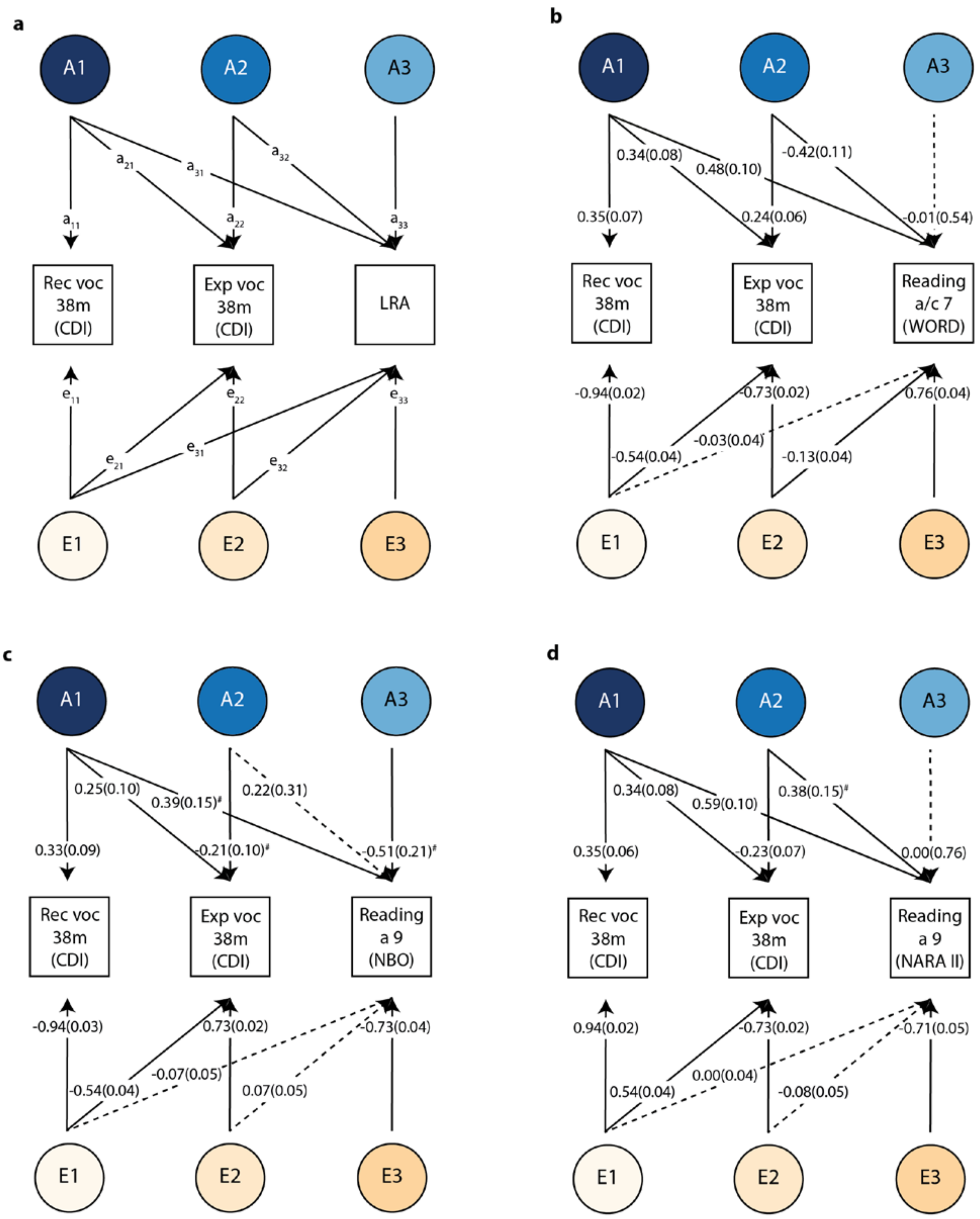
a
A1
A2
A3
a11
a21
a31
a22
a32
a33
Rec voc 38m (CDI)
Exp voc 38m (CDI)
LRA
e11
e21
e31
e22
e32
e33
E1
E2
E3
b
A1
A2
A3
0.34(0.08)
0.48(0.10)
-0.42(0.11)
0.35(0.07)
0.24(0.06)
-0.01(0.54)
Rec voc 38m (CDI)
Exp voc 38m (CDI)
Reading a/c 7 (WORD)
-0.94(0.02)
-0.73(0.02)
0.76(0.04)
-0.03(0.04)
-0.54(0.04)
-0.13(0.04)
E1
E2
E3
c
A1
A2
A3
0.25(0.10)
0.39(0.15)#
0.22(0.31)
0.33(0.09)
0.21(0.10)#
-0.51(0.21)#
Rec voc 38m (CDI)
Exp voc 38m (CDI)
Reading a 9 (NBO)
-0.94(0.03)
0.73(0.02)
-0.73(0.04)
-0.07(0.05)
-0.54(0.04)
0.07(0.05)
E1
E2
E3
d
A1
A2
A3
0.34(0.08)
0.59(0.10)
0.38(0.15)#
0.35(0.06)
-0.23(0.07)
0.00(0.76)
Rec voc 38m (CDI)
Exp voc 38m (CDI)
Reading a 9 (NARA II)
0.94(0.02)
-0.73(0.02)
-0.71(0.05)
0.00(0.04)
0.54(0.04)
-0.08(0.05)
E1
E2
E3

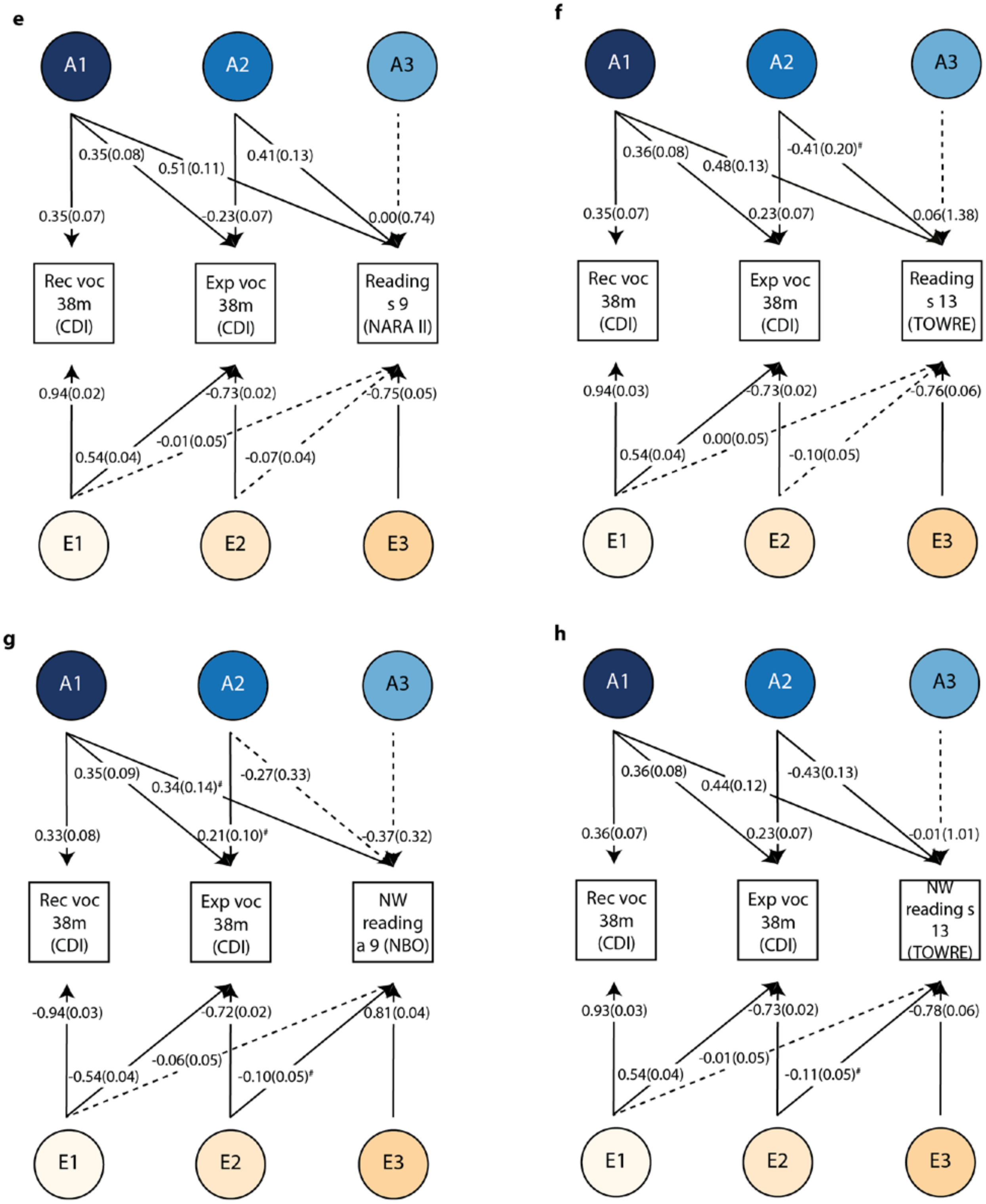
e
A1
A2
A3
0.35(0.08)
0.51(0.11)
0.41(0.13)
0.35(0.07)
-0.23(0.07)
0.00(0.74)
Rec voc 38m (CDI)
Exp voc 38m (CDI)
Reading s 9 (NARA II)
0.94(0.02)
-0.73(0.02)
-0.75(0.05)
0.54(0.04)
-0.01(0.05)
-0.07(0.04)
E1
E2
E3
f
A1
A2
A3
0.36(0.08)
0.48(0.13)
-0.41(0.20)#
0.35(0.07)
0.23(0.07)
0.06(1.38)
Rec voc 38m (CDI)
Exp voc 38m (CDI)
Reading s 13 (TOWRE)
0.94(0.03)
-0.73(0.02)
-0.76(0.06)
0.54(0.04)
0.00(0.05)
-0.10(0.05)
E1
E2
E3
g
A1
A2
A3
0.35(0.09)
0.34(0.14)#
-0.27(0.33)
0.33(0.08)
0.21(0.10)#
-0.37(0.32)
Rec voc 38m (CDI)
Exp voc 38m (CDI)
NW reading a 9 (NBO)
-0.94(0.03)
-0.72(0.02)
0.81(0.04)
-0.54(0.04)
-0.06(0.05)
-0.10(0.05)#
E1
E2
E3
h
A1
A2
A3
0.36(0.08)
0.44(0.12)
-0.43(0.13)
0.36(0.07)
0.23(0.07)
-0.01(1.01)
Rec voc 38m (CDI)
Exp voc 38m (CDI)
NW reading s 13 (TOWRE)
0.93(0.03)
-0.73(0.02)
-0.78(0.06)
0.54(0.04)
-0.01(0.05)
-0.11(0.05)#
E1
E2
E3

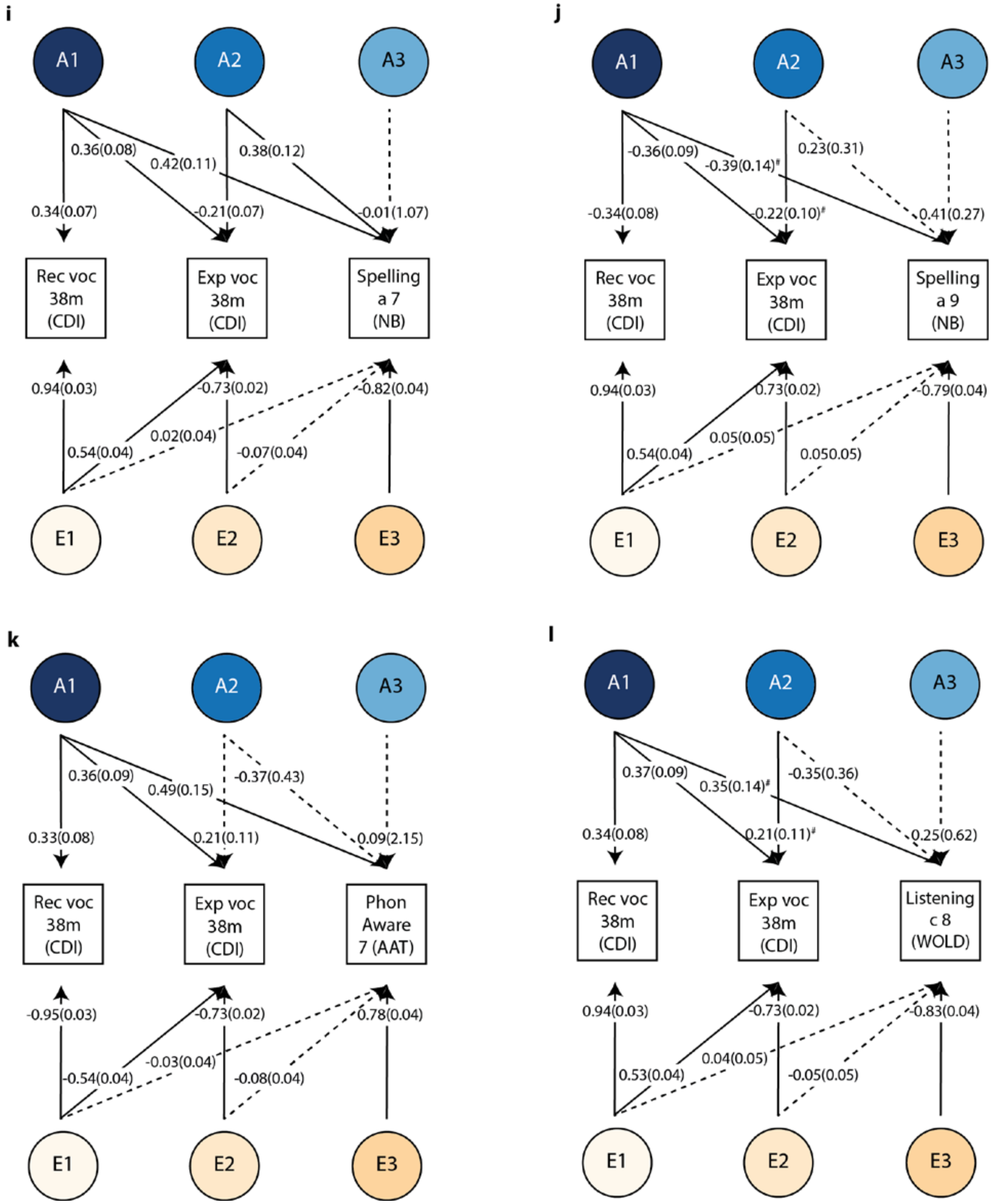
i
A1
A2
A3
0.36(0.08)
0.42(0.11)
0.38(0.12)
0.34(0.07)
-0.21(0.07)
-0.01(1.07)
Rec voc 38m (CDI)
Exp voc 38m (CDI)
Spelling a 7 (NB)
0.94(0.03)
-0.73(0.02)
-0.82(0.04)
0.02(0.04)
0.54(0.04)
-0.07(0.04)
E1
E2
E3
j
A1
A2
A3
-0.36(0.09)
-0.39(0.14)#
0.23(0.31)
-0.34(0.08)
-0.22(0.10)#
0.41(0.27)
Rec voc 38m (CDI)
Exp voc 38m (CDI)
Spelling a 9 (NB)
0.94(0.03)
0.73(0.02)
-0.79(0.04)
0.05(0.05)
0.54(0.04)
0.050.05)
E1
E2
E3
k
A1
A2
A3
0.36(0.09)
0.49(0.15)
-0.37(0.43)
0.33(0.08)
0.21(0.11)
0.09(2.15)
Rec voc 38m (CDI)
Exp voc 38m (CDI)
Phon Aware 7 (AAT)
-0.95(0.03)
-0.73(0.02)
0.78(0.04)
-0.03(0.04)
-0.54(0.04)
-0.08(0.04)
E1
E2
E3
l
A1
A2
A3
0.37(0.09)
0.35(0.14)#
-0.35(0.36)
0.34(0.08)
0.21(0.11)#
0.25(0.62)
Rec voc 38m (CDI)
Exp voc 38m (CDI)
Listening c 8 (WOLD)
0.94(0.03)
-0.73(0.02)
-0.83(0.04)
0.04(0.05)
0.53(0.04)
-0.05(0.05)
E1
E2
E3

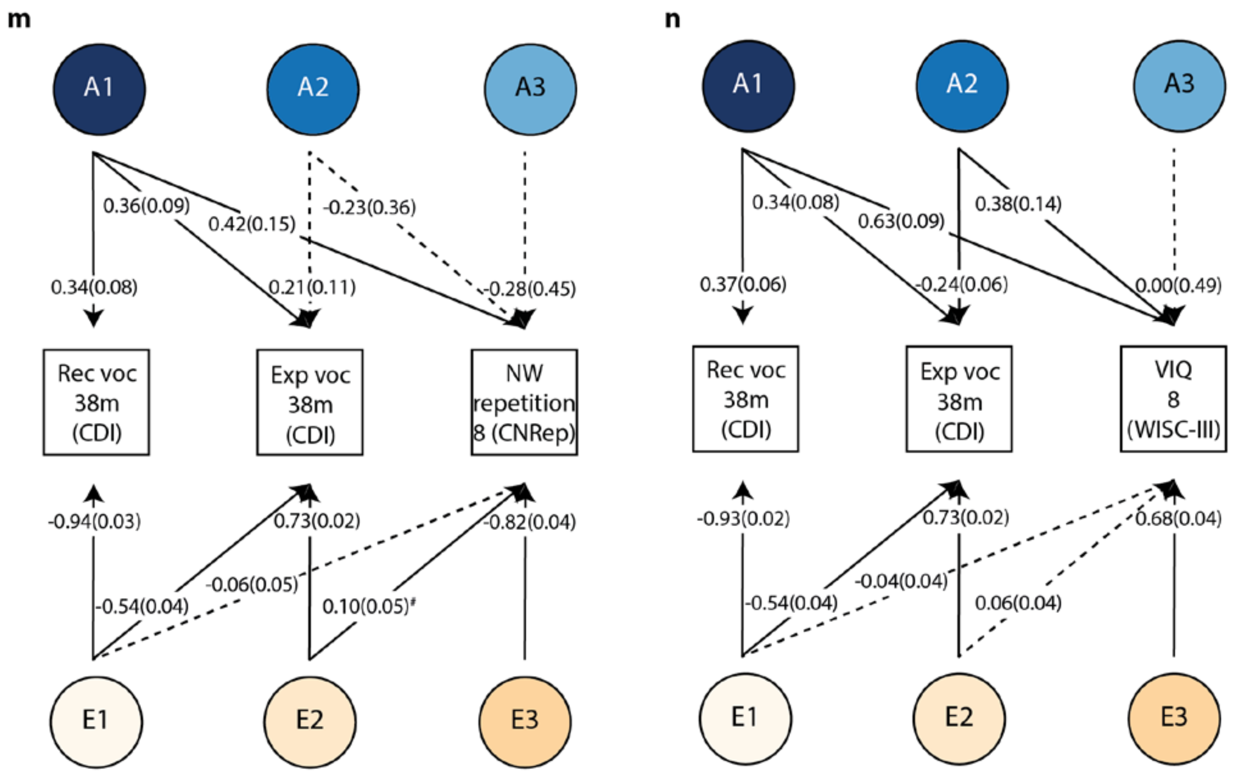

**Supplementary Figure 6: Path models of early vocabulary and mid-childhood to adolescence literacy- and language-related abilities (reverse GSEM)**

Abbreviations: a, accuracy; AAT, Auditory Analysis Test; c, comprehension; CDI, Communicative Development Inventory; CNRep, Children's Test of Nonword Repetition; Exp, expressive; LRA, language- and literacy-related ability; m, months; NARA II, The Neale Analysis of Reading Ability- Second Revised British Edition; NB, ALSPAC-specific assessment developed by Nunes and Bryant; NBO, ALSPAC-specific assessment developed by Nunes, Bryant and Olson; NW, nonword; PhonAware, phonemic awareness; Rec, receptive; s, speed; TOWRE, Test Of Word Reading Efficiency; VIQ, verbal intelligence quotient; voc, vocabulary; WISC-III, Wechsler Intelligence Scale for Children III; WOLD, Wechsler Objective Language Dimensions; WORD, Wechsler Objective Reading Dimension

### Path coefficient passing nominal significance ($P \leq 0.05$), but not the experiment-wide significance threshold ($P \leq 0.005$).

Cholesky decompositions were fitted using GSEM, according to reverse GSEMs and based on all available observations for children across development (N≤6,092). **(a)** Schematic path model with path coefficient labels for a Cholesky decomposition model of vocabulary at 38 months, including receptive and expressive vocabulary (in that order), and one later LRA. **(b-l)** Path models of standardised path coefficients and corresponding standard errors for 13 reverse GSEM models, one for each fitted LRA. Solid lines indicate path coefficients passing a *P*-value threshold of $P \leq 0.05$, dashed lines indicate non-significant path coefficients $P > 0.05$.

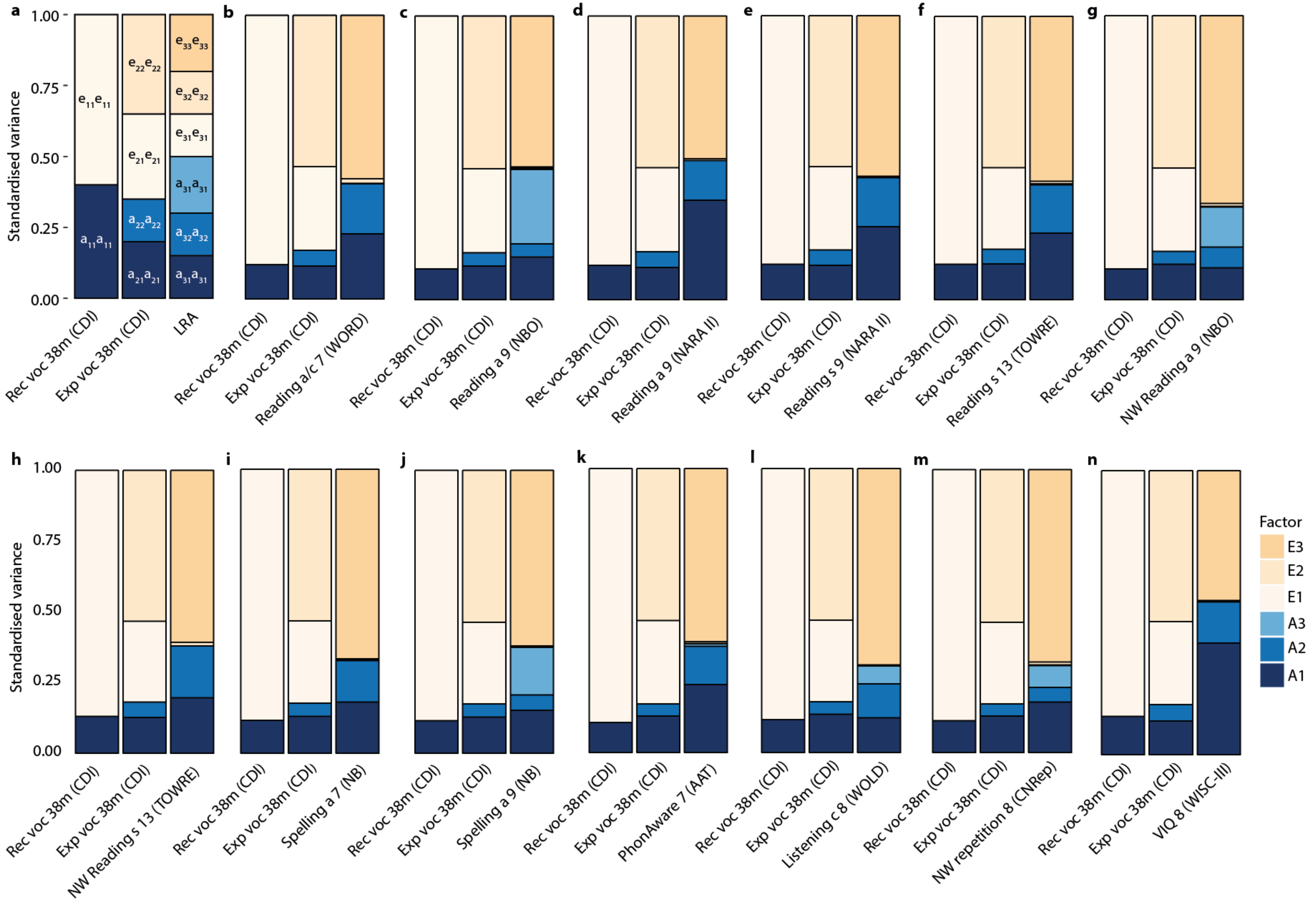

a
b
c
d
e
f
g
h
i
j
k
l
m
n
Standardised variance
1.00
0.75
0.50
0.25
0.00
e11e11
a11a11
e22e22
e21e21
a22a22
a21a21
e33e33
e32e32
e31e31
a31a31
a32a32
a31a31
Rec voc 38m (CDI)
Exp voc 38m (CDI)
LRA
Reading a/c 7 (WORD)
Reading a 9 (NBO)
Reading a 9 (NARA II)
Reading s 9 (NARA II)
Reading s 13 (TOWRE)
NW Reading a 9 (NBO)
NW Reading s 13 (TOWRE)
Spelling a 7 (NB)
Spelling a 9 (NB)
PhonAware 7 (AAT)
Listening c 8 (WOLD)
NW repetition 8 (CNRep)
VIQ 8 (WISC-III)
Factor
E3
E2
E1
A3
A2
A1

Supplementary Figure 7: Variance plots for path models of early vocabulary and mid-childhood to adolescence literacy- and language-related abilities (reverse GSEM)

Abbreviations: a, accuracy; AAT, Auditory Analysis Test; c, comprehension; CDI, Communicative Development Inventory; CNRep, Children's Test of Nonword Repetition; Exp, expressive; LRA, language- and literacy-related ability; m, months; NARA II, The Neale Analysis of Reading Ability- Second Revised British Edition; NB, ALSPAC-specific assessment developed by Nunes and Bryant; NBO, ALSPAC-specific assessment developed by Nunes, Bryant and Olson; NW, nonword; PhonAware, phonemic awareness; Rec, receptive; s, speed; TOWRE, Test Of Word Reading Efficiency; VIQ, verbal intelligence quotient; voc, vocabulary; WISC-III, Wechsler Intelligence Scale for Children III; WOLD, Wechsler Objective Language Dimensions; WORD, Wechsler Objective Reading Dimension

Standardised variance explained by genetic and residual factors as derived by Cholesky decompositions using reverse GSEM (Supplementary Figure 6), based on all available observations for children across development (N≤6,092). **(a)** Variance plot with path coefficient labels for a Cholesky decomposition model of vocabulary at 38 months, including receptive and expressive vocabulary (in that order), and one later LRA. **(b-l)** Standardised variance explained by genetic and residual factors as modelled in 13 reverse GSEM models, one for each fitted LRA.

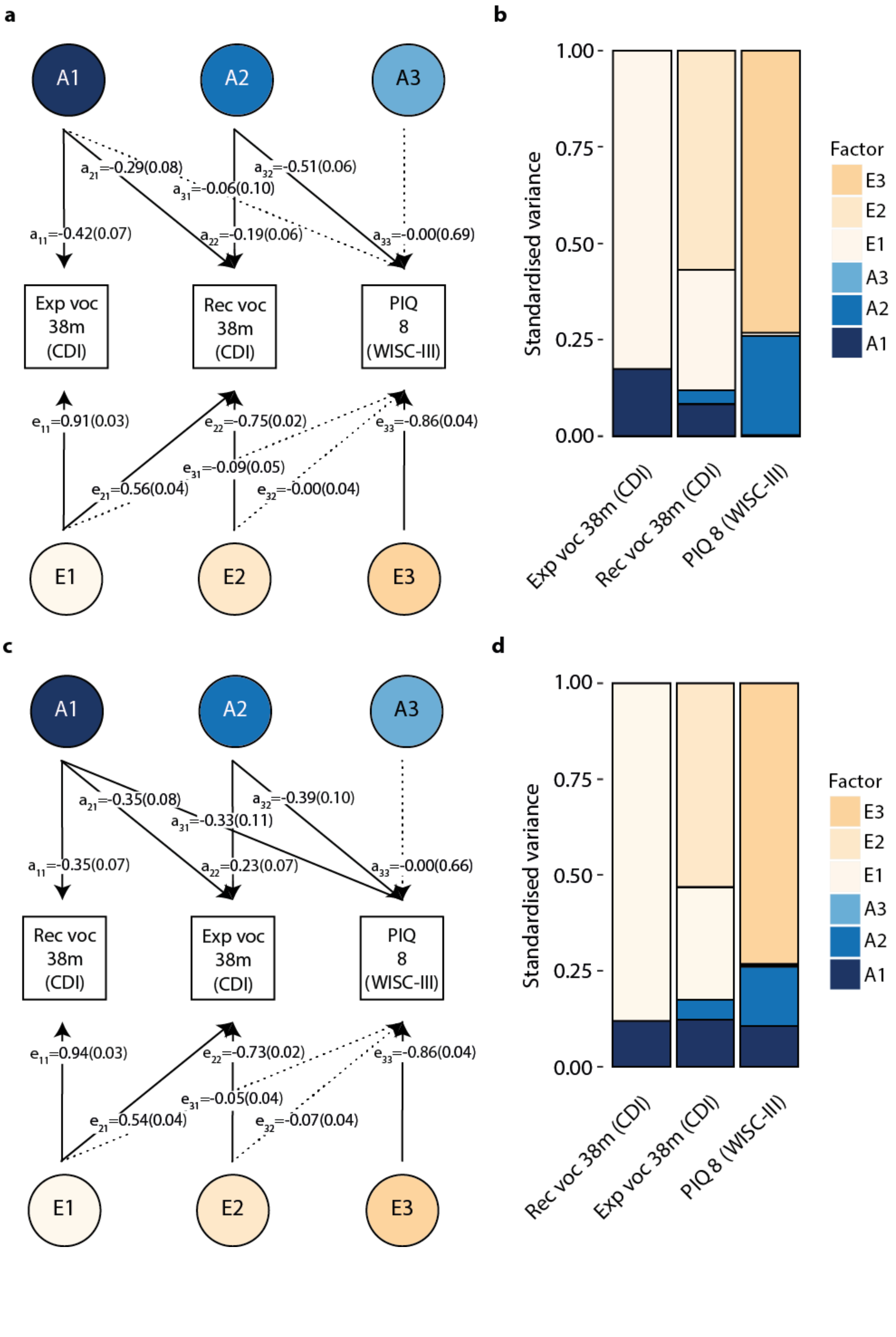

a
A1
A2
A3
a21=-0.29(0.08)
a32=-0.51(0.06)
a31=-0.06(0.10)
a11=-0.42(0.07)
a22=-0.19(0.06)
a33=-0.00(0.69)
Exp voc 38m (CDI)
Rec voc 38m (CDI)
PIQ 8 (WISC-III)
e11=0.91(0.03)
e22=-0.75(0.02)
e33=-0.86(0.04)
e31=-0.09(0.05)
e21=0.56(0.04)
e32=-0.00(0.04)
E1
E2
E3
b
Standardised variance
1.00
0.75
0.50
0.25
0.00
Exp voc 38m (CDI)
Rec voc 38m (CDI)
PIQ 8 (WISC-III)
Factor
E3
E2
E1
A3
A2
A1
c
A1
A2
A3
a21=-0.35(0.08)
a32=-0.39(0.10)
a31=-0.33(0.11)
a11=-0.35(0.07)
a22=0.23(0.07)
a33=-0.00(0.66)
Rec voc 38m (CDI)
Exp voc 38m (CDI)
PIQ 8 (WISC-III)
e11=0.94(0.03)
e22=-0.73(0.02)
e33=-0.86(0.04)
e31=-0.05(0.04)
e21=0.54(0.04)
e32=-0.07(0.04)
E1
E2
E3
d
Standardised variance
1.00
0.75
0.50
0.25
0.00
Rec voc 38m (CDI)
Exp voc 38m (CDI)
PIQ 8 (WISC-III)
Factor
E3
E2
E1
A3
A2
A1

Supplementary Figure 8: Path model and variance plot for early vocabulary and mid-childhood performance intelligence scores

Abbreviations: CDI, Communicative Development Inventory; Exp, expressive; m, months; Rec, receptive; voc, vocabulary; PIQ, performance intelligence quotient; WISC-III, Wechsler Intelligence Scale for Children III

A Cholesky decomposition was fitted using GSEM, according to **(a,b)** forward GSEM and **(c,d)** reverse GSEM, based on all available observations for children across development (N≤6,092). **(a)** Path model of standardised path coefficients and corresponding standard errors for a Cholesky decomposition of vocabulary at 38 months, including expressive and receptive vocabulary (in that order), and performance intelligence scores at 8 years. Solid lines indicate path coefficients passing a *P*-value threshold of *P*≤0.05, dashed lines indicate non-significant path coefficients (*P*>0.05). **b)** Standardised variance explained by genetic and residual factors modelled in a. **(c)** Path model of standardised path coefficients and corresponding standard errors for a Cholesky decomposition of vocabulary at 38 months, including receptive and expressive vocabulary (in that order), and performance intelligence scores at 8 years. Solid lines indicate path coefficients passing a *P*-value threshold of *P*≤0.05, dashed lines indicate non-significant path coefficients (*P*>0.05). **d)** Standardised variance explained by genetic and residual factors modelled in c.
